# A window trial in metastatic pancreatic ductal adenocarcinoma reveals resistance mechanisms to targeting the KRAS-MEK pathway

**DOI:** 10.64898/2025.12.12.693851

**Authors:** Motoyuki Tsuda, Dove Keith, Colin J. Daniel, Carl Pelz, Katie E. Blise, Tugba Y. Ozmen, Furkan Ozmen, Kevin Hawthone, Jennifer R. Eng, Xi Li, Xiaoyan Wang, Hayley Zimny, Shamilene Sivagnanam, Konjit Betre, Nell Kirchberger, Benson Chong, Jinho Lee, Aubry Matter, Alexander Smith, Ethan S. Agritelley, Trent Waugh, Ashwin Sannecy, Kimberly Alarcon, Isaac Youm, Sara Protzek, Julian Egger, Isabel A. English, Min Yang, Vidhi M. Shah, Jason M. Link, Allison L. Creason, Patrick J. Worth, Shaun M. Goodyear, Koei Chin, John L. Muschler, Christopher G. Suciu, Christopher L. Corless, Sha Cao, Laura Soucek, Adel Kardosh, Lisa M. Coussens, Jonathan R. Brody, Charles D. Lopez, Gordon B. Mills, Rosalie C. Sears

## Abstract

Copy number alterations of *KRAS,* mutated in over 90% of pancreatic ductal adenocarcinomas (PDAC), and *MYC* occur in 30-40% of PDAC. Here we demonstrate that *KRAS* and *MYC* are frequently co-gained and accompanied with worse prognosis in PDAC. In a Window-of-Opportunity clinical trial for metastatic PDAC, serial biopsies and deep multi-omics analyses were utilized to explore resistance mechanisms to MEK inhibition, as a surrogate for KRAS inhibition. Tumors from four of 14 patients showed Ki-67/CA19-9-based biomarker response (BR). Non-BR tumors were enriched for *KRAS*/*MYC* co-gain and *KRAS^G12D^* variant. A transcriptomic signature of BR tumors was inversely correlated with *KRAS^G12D^*/*MYC* co-gain in a large PDAC dataset and predictive for KRAS inhibitor response in multiple models. Finally, co-targeting KRAS and MYC was synergistic in *KRAS^G12D^*/*MYC* co-gain PDAC. Together, this study provides insight into KRAS inhibitor resistance and supports MYC as an important target to improve patient outcomes in this deadly disease.

## Introduction

Pancreatic Cancer is the third leading cause of cancer-related deaths in the United States^1^. Pancreatic ductal adenocarcinoma (PDAC), the most common form of the disease, the 5-year survival rate is only 8%. Surgical intervention is possible in fewer than 20% of patients, and survival rates for patients with metastatic PDAC is a dismal 3%. Such dismal outcomes underscore the urgent need for new systemic treatments to improve outcomes for patients with PDAC. Commonly referred to as the ‘Big 4’, *KRAS*, *CDKN2A*, *TP53*, and *SMAD4,* are key gene mutations associated with driving PDAC^2,3^. Of these, *KRAS* mutations occur in over 90%, and the gene is amplified in ∼40% of PDAC cases; reinforcing its role as a dominant oncogenic driver^4^. Mechanistically, the KRAS-MEK-ERK signaling cascade is a crucial downstream mediator of mutant KRAS^5,6^. While the addition of a MEK inhibitor to standard chemotherapy in clinical trials has not yielded superiority over standard chemotherapy, subgroup analyses suggested that *KRAS* wild-type and *KRAS^G12R^* patients may experience more favorable responses to MEK inhibition^7–9^, though the mechanisms underlying response and resistance remain incompletely understood. The emergence of KRAS targeted inhibitors in clinical trials emphasizes the significance of elucidating mechanisms governing resistance and responsiveness to targeting KRAS signaling, including MEK inhibition—a critical downstream effector of KRAS. Understanding these mechanisms is likely to inform the future success of KRAS-targeted therapy in PDAC^10,11^.

Biopsy-rich window-of-opportunity (WOO) clinical trials can provide detailed information on molecular and cellular responses to targeted therapeutics^12^. At the OHSU Knight Cancer Institute, we have implemented a highly adaptable, multi-arm WOO clinical trial platform to assess the biological activity of investigational agents over a 10-day treatment period in patients with metastatic pancreatic cancer (NCT04005690)^13^. Utilizing this window approach, paired pre-and post-treatment biopsies are subjected to multi-omic analyses, including DNA-seq, RNA-seq (bulk and single-cell [sc]), spatial multiplex immunohistochemistry (mIHC), Clinical Laboratory Improvement Amendments (CLIA) IHC, cyclic immunofluorescence (cycIF), and NanoString digital spatial profiling (DSP). Furthermore, patient-derived models, including patient-derived cell lines (PDCLs) and patient-derived xenografts (PDXs), are generated to assess long-term drug response and validate key findings.

In this study, we took advantage of both a large DNA-seq and RNA-seq dataset of over 300 PDACs^3^ and our WOO trial platform data to investigate the role of the KRAS-MEK-ERK signaling pathway and its downstream effector, the c-MYC (MYC) oncoprotein in PDAC metastasis and therapeutic resistance. We found that *KRAS*/*MYC* gene co-gain is common in PDAC, most prominently in metastases, along with marked upregulation of KRAS, ERK, and MYC activity signatures. We utilized the WOO trial platform to clinically investigate targeting the KRAS-MEK pathway with the FDA-approved MEK inhibitor, cobimetinib (Genentech, CA), in patients with metastatic PDAC^13^, and identified biomarker-responsive (BR) tumors based on clinical serum glycan carbohydrate antigen 19-9 (CA19-9) and tissue Ki-67 responses. Multi-omic analyses revealed molecular and cellular features in the tumor and microenvironment associated with MEK inhibitor sensitivity and resistance in metastatic PDAC. Notably, we found that *KRAS^G12D^* mutation and *KRAS^G12D^*/*MYC* co-gain are resistance mechanisms to MEK inhibition and that this mechanism was transferable to KRAS inhibition utilizing *in vitro* and *in vivo* patient-derived models and a genetic PDAC mouse model. Further, we found the concurrent inhibition of KRAS and MYC is synergistic in *KRAS^G12D^*/*MYC* co-gain PDAC. Overall, our study provides new insights into resistance mechanisms associated with targeting KRAS-MEK signaling that are translatable to ongoing KRAS inhibitor trials, as well as key to informing the optimization of targeted therapeutic strategies that can advance precision medicine approaches in PDAC.

## Results

### *KRAS* and *MYC* co-gain and signaling pathway co-activation are enriched in aggressive liver metastatic and poor prognosis PDAC

Aside from *KRAS* being nearly ubiquitously mutated in PDAC, the oncogene also exhibits copy number changes as loss of heterozygosity of the wild-type allele and/or gain in mutant and/or WT alleles in disease progression^4^. A large-scale genomic study showed that *KRAS* gain confers a survival disadvantage^14^, however corresponding transcriptomic changes underlying the biology have not been fully elucidated. Taking advantage of our center’s PDAC dataset (Brenden-Colson Center for Pancreatic Care (BCCPC) dataset; n=308 for copy number variation (CNV) data, n=310 for RNA-seq data, and n=286 with both CNV and RNA-seq), we investigated transcriptomic change by *KRAS* CNVs. The dataset was divided into five CNV types: *KRAS* Normal/Loss (2 copies or 1 copy of wild type, n=203), *KRAS* LOH (loss of heterozygosity, 1 copy of mutant, n=13), *KRAS* CN-LOH (copy neutral LOH, 2 mutant copies, n=7), *KRAS* minor gain (3 copies, n=60) and *KRAS* major gain (≧4 copies, n=25). *KRAS* LOH, minor gain and major gain trended toward worse survival in our dataset compared to *KRAS* Normal/Loss but did not reach significance even when combining these genotypes as *KRAS* imbalance (Figure S1A). Gene Set Variation Analysis (GSVA) of MSigDB Hallmark gene sets plus the PDAC KRAS/ERK gene set^15^ showed the *KRAS* LOH, minor gain and major gain cohorts possessed higher KRAS/ERK signaling activity, cell cycle and replication stress (RS)-related E2F and G2/M checkpoint, and MYC target gene signatures, indicating that *KRAS* CNVs significantly increase downstream effector pathways over *KRAS* copy number Normal/Loss (Figure S1B). Since *KRAS* minor gain and major gain have similar pathway activation, hereafter they were combined as *KRAS* gain. Combining LOH and CN-LOH as LOH indicated some distinct pathway activation compared with *KRAS* gain and *KRAS* Normal/Loss including TGFb, NFkB, and NOTCH signaling (Figure S1C).

*KRAS* regulates MYC protein stability and activity via ERK-mediated phosphorylation^5,16–18^. As a well-established master oncogene, *MYC* is often gained in PDAC (27%-48%)^3,19–21^. High-level amplification of *MYC* is associated with poor survival in PDAC^20^, however, the biological impact of low-level *MYC* gain is less characterized. The dataset was divided into 3 CNV types: MYC Normal/Loss (n=205), MYC minor gain (3 copies, n=72), and MYC major gain (≧4 copies, n=31). By analyzing RNA-seq data, we found that, similar to *KRAS* gain, both *MYC* minor gain and major gain exhibited higher activation of KRAS/ERK, cell cycle and RS pathways, and MYC signaling (Figure S1D), although the difference of minor gain or major gain did not significantly impact patient survival in our BCCPC dataset (Figure S1E). Given the KRAS and MYC relationship and their frequent gain rates, we hypothesized that *KRAS* and *MYC* gene co-gains may occur in PDAC, leading to cooperative signaling activity. To test this, we analyzed *KRAS* and *MYC* copy number alterations in our PDAC dataset and observed significant enrichment of *MYC* gain in *KRAS* gain (FDR=0.005) compared with *KRAS* Normal/Loss (Figure 1A). This finding was validated in three independent external datasets (TCGA: p=0.0002, CPTAC: p=0.0006, UTSW: p=0.0143) (Figures 1A and S2A). Further analysis revealed that metastases are enriched in *KRAS*/*MYC* co-gain compared with their copy number Normal/Loss (FDR=0.01) (Figure 1B).

**Figure 1:**
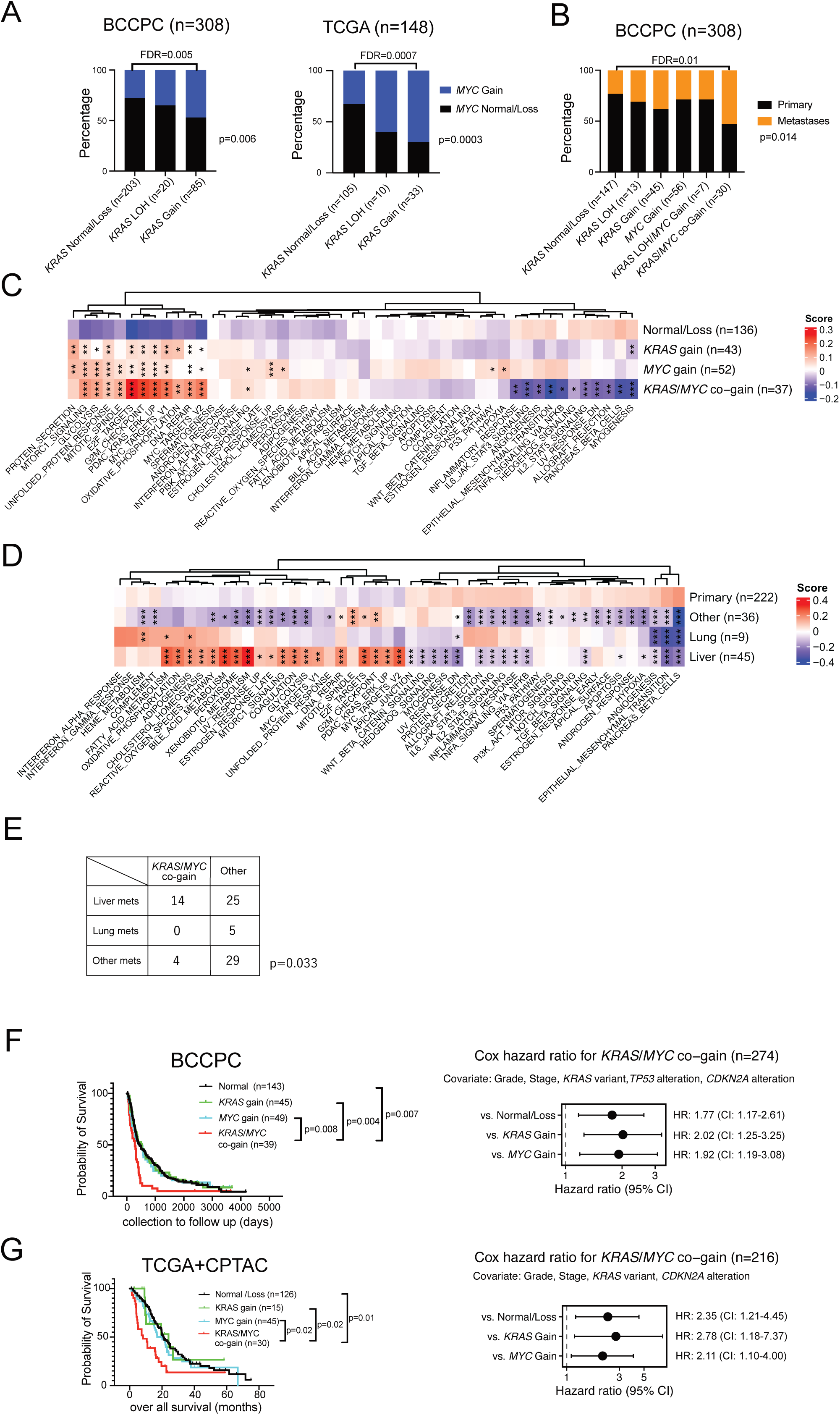
*KRAS* and *MYC* gene co-gain is enriched in liver metastatic PDAC and predicts poor prognosis. (**A**) Relationship between *KRAS* copy number variation (CNV) and *MYC* CNV in the Brenden-Colson Center for Pancreatic Care (BCCPC) cohort (left; n=308 with CNV data) and TCGA cohort (right; n=148 with CNV data). Normal/Loss includes CN-neutral and CN=1 without mutation. Gain includes CN≥3. LOH indicates *KRAS* mutant samples with loss of heterozygosity in CN=1 or 2. Whole group comparison was evaluated by Fisher’s exact test. Pairwise comparisons were evaluated by Wilcoxon test and corrected by FDR. (**B**) The relationship between *KRAS* and *MYC* CNV status and metastases. BCCPC cohort. Whole group comparison was evaluated by Fisher’s exact test. Pairwise comparisons were evaluated by Wilcoxon test and corrected by FDR. (**C**) Heatmap of mean GSVA scores of MSigDB Hallmark and KRAS/ERK signatures across *KRAS* and *MYC* copy number gain status. BCCPC cohort (n=268, with CNV and RNA-seq data, *KRAS* LOH was excluded.) One-way ANOVA with post-hoc Dunnet’s test (Ref: Normal/Loss). *: p<0.05, **: p<0.01, ***: p<0.001. (**D**) Heatmap of mean GSVA scores of Hallmark and KRAS/ERK signatures in metastasis and primary PDAC. BCCPC cohort (n=312; with RNA-seq data), One-way ANOVA with post-hoc Dunnet’s test (Ref: Primary). *: p<0.05, **: p<0.01, ***: p<0.001. (**E**) The numbers of metastases divided by metastatic sites. Compared between *KRAS*/*MYC* copy number co-gain and other groups. BCCPC cohort. *KRAS* LOH was excluded. Fisher’s exact test. (**F**) Kaplan-Meier (KM) survival plot and Forest Plot regarding *KRAS* and *MYC* gain status in BCCPC cohort (n=276; with CNV data, *KRAS* LOH and patients died within 30 days of enrollment were excluded.). Normal/Loss/Gain only. Multivariate analysis adjusted by significant factors in Univariate analysis (tumor stage, tumor grade, *KRAS* mutant variant, and *TP53* and *CDKN2A* mutation status). P value is from Cox Hazard Multivariate analysis. (**G**) KM survival plot and Forest Plot regarding *KRAS* and *MYC* gain status in CPTAC+TCGA cohort(n=216; with CNV data, *KRAS* LOH and patients died within 30 days of enrollment were excluded.). Normal/Loss/Gain only. Multivariate analysis adjusted by significant factors in Univariate analysis (tumor stage, tumor grade, *KRAS* mutant variant and *CDKN2A* mutation status. P value is from Cox Hazard Multivariate analysis.

Transcriptionally, GSVA revealed that *KRAS* gain, *MYC* gain and *KRAS/MYC* co-gain each correlated with marked upregulation of KRAS/ERK, cell cycle/RS pathways, and MYC signatures, with co-gain showing the greatest enrichment in our dataset compared with Normal/Loss (Fig. 1C). While MYC is known to be downstream of the KRAS/ERK pathway, *MYC* gain alone was associated with KRAS/ERK pathway activation in addition to PI3K pathway activation. ERBB signaling upstream of KRAS is a known amplifier of KRAS signaling^22,23^, and *MYC* copy gain was associated with increased expression of *ERBB2/HER2* (FDR=0.011), *ERBB3/HER3* (FDR=0.026) and *KRAS* (FDR=0.073), while EGFR was not increased (FDR=0.57) (Figure 2B). Further analysis of inferred protein activity using Virtual Inference of Protein Activity by Enriched Regulon (VIPER) analysis^3^ showed upregulation of ERBB2 (FDR=0.000034), ERBB3 (FDR=0.0019), KRAS (FDR=0.00019), and EGFR (FDR=0.018) protein activity in *MYC* gain only compared to Normal/Loss (Figure S2C). Together with our previous finding that MYC deregulation increases phosphorylated ERBB2 expression in an *in vivo* breast cancer model^24^, these data suggest that *KRAS* and *MYC* copy number gains coordinately enhance KRAS/ERK and MYC signaling activity potentially through a positive feedback loop involving ERBB2 activation.

**Figure 2:**
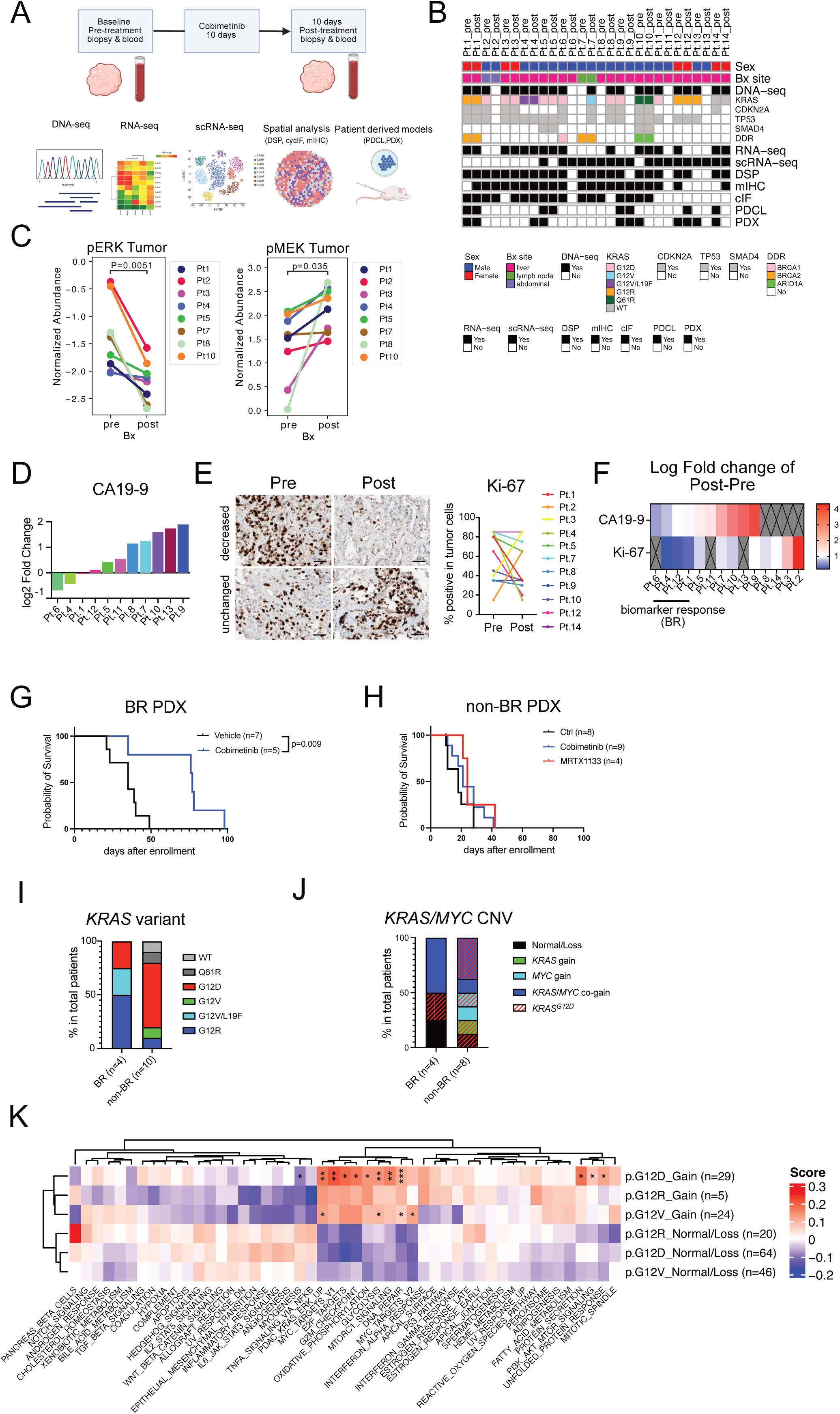
WOO trial design and features of biomarker response related to *KRAS^G12D^*/*MYC* co-gain. (**A**)Schematic of WOOM trial. (**B**) Summary of Patients and obtained samples. (n=14 patients, 28 biopsies total). (**C**) Digital Spatial Profiling (DSP). Comparison of normalized protein abundance between pre- and post-treatment. Phospho-ERK (Left: n=8) and phospho-MEK (Right: n=8). (**D**) Log2 fold change of serum CA19-9 between start-date and end-date of 10-day treatment (n=11). (**E**) Change of Ki-67 staining positivity between the first and second biopsies (n=11). (**F**) Heatmap showing changes in CA19-9 and Ki-67. (**G, H**) Survival plot of cobimetinib or MRTX1133 treated patient-derived xenograft (PDX). (**G**) Biomarker Response (BR) PDX (Ctrl: n=7, Cobimetinib: n=5) (**H**) non-BR PDX (Ctrl: n=8, Cobimetinib: n=9, MRTX1133: n=4). **(I**) *KRAS* mutation variants in BR (n=4) and non-BR (n=10) patients. (**J**) *KRAS*/*MYC* CNV status of BR (n=4) and non-BR (n=8) patients. *KRAS^G12D^* patients (n=1 in BR and n=6 in nonBR) are shading in red. (**K**) Heatmap of mean GSVA scores of Hallmark and KRAS/ERK signatures regarding *KRAS* mutation variants and gain. BCCPC cohort (n=211; *KRAS^G12D^, KRAS^G12V^*, *KRAS^G12R^* only). One-way ANOVA with post-hoc Dunnet’s test (ref: G12V Normal/Loss). *: p<0.05, **: p<0.01, ***: p<0.001.

Separating primary tumors from metastases indicated that *KRAS*/*MYC* co-gain in primary PDAC associates with increased KRAS/ERK and MYC signatures compared to *KRAS* or *MYC* single gain or Normal/Loss primary PDAC (Figure S2D). In contrast, *KRAS*/*MYC* co-gain and *KRAS* or *MYC* single gain is associated with activation of KRAS/ERK and MYC pathways in metastatic PDAC, suggesting additional tumor evolution or selection in *KRAS* or *MYC* gain tumors during metastatic development. Liver metastases occur in more than 70% of PDAC metastases and significantly contribute to mortality compared with other metastatic sites^25^. We compared transcriptomic signatures between primary tumors and different metastatic sites with differing prognosis; whereby patients with lung metastasis have increased survival time over patients with liver metastasis^3^ (Primary: n = 222; Liver metastases: n = 45; Lung metastases: n = 9; Other metastases: n = 36). The analysis revealed transcriptomic heterogeneity across metastatic sites with increased KRAS/ERK signatures, MYC signatures, and cell cycle and RS-related E2F and G2M checkpoint signatures specifically in liver metastases (Figure 1D). *KRAS*/*MYC* co-gain was also significantly increased in liver metastases compared to other metastatic sites (Figure 1E). Finally, *KRAS*/*MYC* co-gain was predictive of poorer overall survival, compared with normal *KRAS*/*MYC* copy number and single *KRAS* or *MYC* gain, as demonstrated in the BCCPC and the TCGA+CPTAC datasets (Figures 1G and 1H; Table S1 and S2). This survival disadvantage was also observed when analyzing only primary PDAC in the BCCPC dataset (Figure 2E; Table S3). These findings collectively highlight *KRAS*/*MYC* co-gain and cooperative KRAS-MEK-ERK and MYC pathway signaling association with PDAC liver metastasis and poorer prognosis.

### A window of opportunity trial in metastatic PDAC (WOOM) with the MEK Inhibitor cobimetinib

Given the prevalence of *KRAS*/*MYC* co-gain in poor outcome and liver metastatic PDAC, and the increase in KRAS/MEK/ERK signaling activity, we aimed to assess the efficacy of targeting this pathway in metastatic patients utilizing our WOO PDAC trial platform to evaluate the FDA-approved MEK inhibitor cobimetinib (60 mg PO QD for 10 days). The WOOM trial design included deep omic profiling of pre- and post-treatment biopsies immediately flanking a ten-day treatment period with the goal of providing rich data on how tumors evolve under therapeutic stress (Figure 2A). Of the 14 patients enrolled to the cobimetinib arm, 12 had metastatic disease to the liver. DNA-seq, RNA-seq, scRNA-seq, cycIF, mIHC, and DSP analyses were successfully conducted in the majority of patients (Figure 2B). While the generation of patient-derived models was challenging due to the small size of each biopsy; 5 patient-derived cell lines (PDCLs) and 9 patient-derived xenograft (PDX) models were successfully established.

To evaluate the on-target activity of cobimetinib in patient tumors, we assessed ERK phosphorylation (pERK) using the CLIA DSP platform and observed its downregulation in both tumor and tumor-adjacent stromal regions across all post-treatment samples (Figure 2C and S3A). Consistent with previous findings^26^, all patient tumors also exhibited feedback-induced MEK phosphorylation (pMEK); altogether indicating that cobimetinib effectively inhibited the target at the dose and time assessed. To evaluate tumor response to treatment, we measured levels of CA19-9, a clinical blood-based tumor biomarker elevated in ∼75% of PDAC patients^27^. Blood CA19-9 levels were assessed in samples collected on the day of each biopsy. A total of 11 of the 14 patients had CA19-9 elevation, 2 patients showed a ≥15% decrease, and two others showed stable levels (Figure 2D). Next, we assessed tumor proliferation via CLIA Ki-67 IHC staining, in which a strong reduction (i.e., ≥50% decrease in Ki-67 expression) was noted in 3 of 11 paired specimens (Figure 2E). For one patient with decreased tumor Ki-67, CA19-9 was also decreased, and for two other patients with decreased Ki-67, CA19-9 was stable. These three tumors were classified as biomarker responsive (BR). Although Ki-67 change was not clinically available for the patient with the largest decrease in CA19-9 due to lack of tumor cells in the post-treatment biopsy IHC specimen, based on the consistency between CA19-9 and Ki-67 change in the other patients, we also classified this patient’s tumor as BR. Therefore, a total of 4 tumors were classified as BR out of 14 tumors analyzed (Figure 2F).

Since the WOO trial is not a therapeutic trial, any association between BR status and patient long-term response is unclear. Indeed, sample collection to follow-up showed no survival difference in BR patients, although they trended to have longer survival from diagnosis to follow up (Figures S3C and S3D); potentially indicating clinically-relevant biological differences underlying BR and non-BR tumors. Therefore, to address the therapeutic potential of cobimetinib in BR tumors, we treated PDX models derived from both BR and non-BR tumors (Figures 2G and 2H). The BR-PDX model demonstrated prolonged survival, whereas the non-BR-PDX model did not benefit from cobimetinib treatment, suggesting that BR status correlates with therapeutic benefit. Moreover, the KRAS^G12D^ specific inhibitor, MRTX1133, did not improve survival in the non-BR-PDX model harboring *KRAS^G12D^*mutation, suggesting shared resistance mechanisms between MEK and KRAS inhibitors (Figure 2H).

### The WOOM trial identifies *KRAS* variant-specific enrichment and copy number gains in MEK Inhibitor non-biomarker response (non-BR) tumors

To investigate the mechanisms underlying differential biomarker response, we initially analyzed the genomic biopsy data of pre-treatment samples grouped as BR or non-BR. Genomic analysis revealed a trend toward *KRAS^G12R^* enrichment in BR tumors and *KRAS^G12D^* enrichment in non-BR tumors (Figure 2I). This finding aligns with prior clinical trials targeting MEK inhibitors, including cobimetinib^28^. Given that *KRAS*/*MYC* co-gain observed in our large dataset was increased in liver metastatic PDAC, as well as associated with high KRAS/MEK/ERK and MYC pathway signaling, and independently predictive of worse survival, we sought to compare the *KRAS* and *MYC* CNV status along with variant type between BR and non-BR tumors. *KRAS* or *MYC* gain was only observed in non-BR tumors, but *KRAS*/*MYC* co-gain was not different between BR and non-BR tumors (Figure 2J). However, *KRAS^G12R^* and *KRAS^G12V^* mutations were observed in BR tumors with *KRAS*/*MYC* co-gain, while *KRAS^G12D^* mutation with gains in *KRAS*, *MYC* or both were only present in non-BR tumors (Figure 2J). Although none of the genomic analyses here met statistical significance due to the small sample size, the data suggests that *KRAS^G12D^* mutation and *KRAS/MYC* co-gain might cooperatively contribute to cobimetinib non-BR status.

Analysis in our larger BCCPC dataset showed significant survival advantage of *KRAS^G12V^* and *KRAS^Q61H^* compared with *KRAS^G12D^* mutation by multivariable Cox proportional hazards regression analysis (Figure S3D). Together with the recent reports that showed better survival of PDAC patients with *KRAS^G12V^* and *KRAS^G12R^* mutant tumors compared to *KRAS^G12D^* mutant tumors^14,29^, this data supports the different biology of *KRAS^G12V^* and *KRAS^G12R^*mutant tumors compared with *KRAS^G12D^* mutant tumors. Pathway analysis showed that *KRAS^G12D^* mutant tumors have increased activation of the PI3K pathway in addition to KRAS/ERK and MYC pathways, compared with *KRAS^G12V^*and *KRAS^G12R^* mutant, specifically with CN gain (Figure 2K). The lack of PI3K activation associated with *KRAS^G12R^* and *KRAS^G12V^* aligns with previous findings^28,30^. Together with the KRAS-ERK, MYC and PI3K pathway activation seen with *MYC* gain (Figure 1C, and S1D), these findings underscore the importance of KRAS, MYC and PI3K pathway activity in MEK inhibitor resistant PDAC.

### Non-BR pre-treatment tumors show co-activation of KRAS/ERK and MYC pathways and basal-like subtype

We performed Gene Set Enrichment Analysis (GSEA) of RNA-seq data from the pre-treatment biopsies to elucidate transcriptomic differences between BR and non–BR tumors. Significant upregulation of KRAS/ERK and MYC pathways, RS-related G2M checkpoint and E2F target pathways, and Interferon response pathways were seen in non-BR tumors (Figure 3A). scRNA-seq from pre-treatment biopsies showed that the oncogenic bulk RNA-seq signatures were derived from cancer cells, while other signatures like interferon-alpha and interferon-gamma were immune cell dominant (Figure S4A and S4B). To analyze tumor-intrinsic innate mechanisms of resistance in our scRNA-seq data obtained from the pre-treatment biopsies, we integrated a published scRNA-seq PDAC atlas dataset as a reference dataset to improve the accuracy of the clustering (Figure 3B)^31^. Unsupervised clustering showed different compositions of several clusters between BR and non-BR cancer cells. GSEA between BR enriched clusters (Clusters 0, 3 and 5) and non-BR enriched clusters (Clusters 1 and 4) confirmed the differences of pathway activity observed in bulk RNA-seq, particularly cell intrinsic pathways including KRAS/ERK and MYC pathways and RS-related pathways upregulated in non-BR tumors (Figure 3C). Analysis of molecular subtype difference between pre-treatment BR and non-BR biopsies showed non-BR tumors displayed higher basal-like/squamous and lower classical-like subtype signatures compared to BR tumors (Figure 3D). Sample-level score of scRNA-seq subtype signatures also indicated that BR tumors had significantly higher sc-Classical signature scores and tended to have lower sc-Basal signature scores (Figure 3E)^32^.

**Figure 3:**
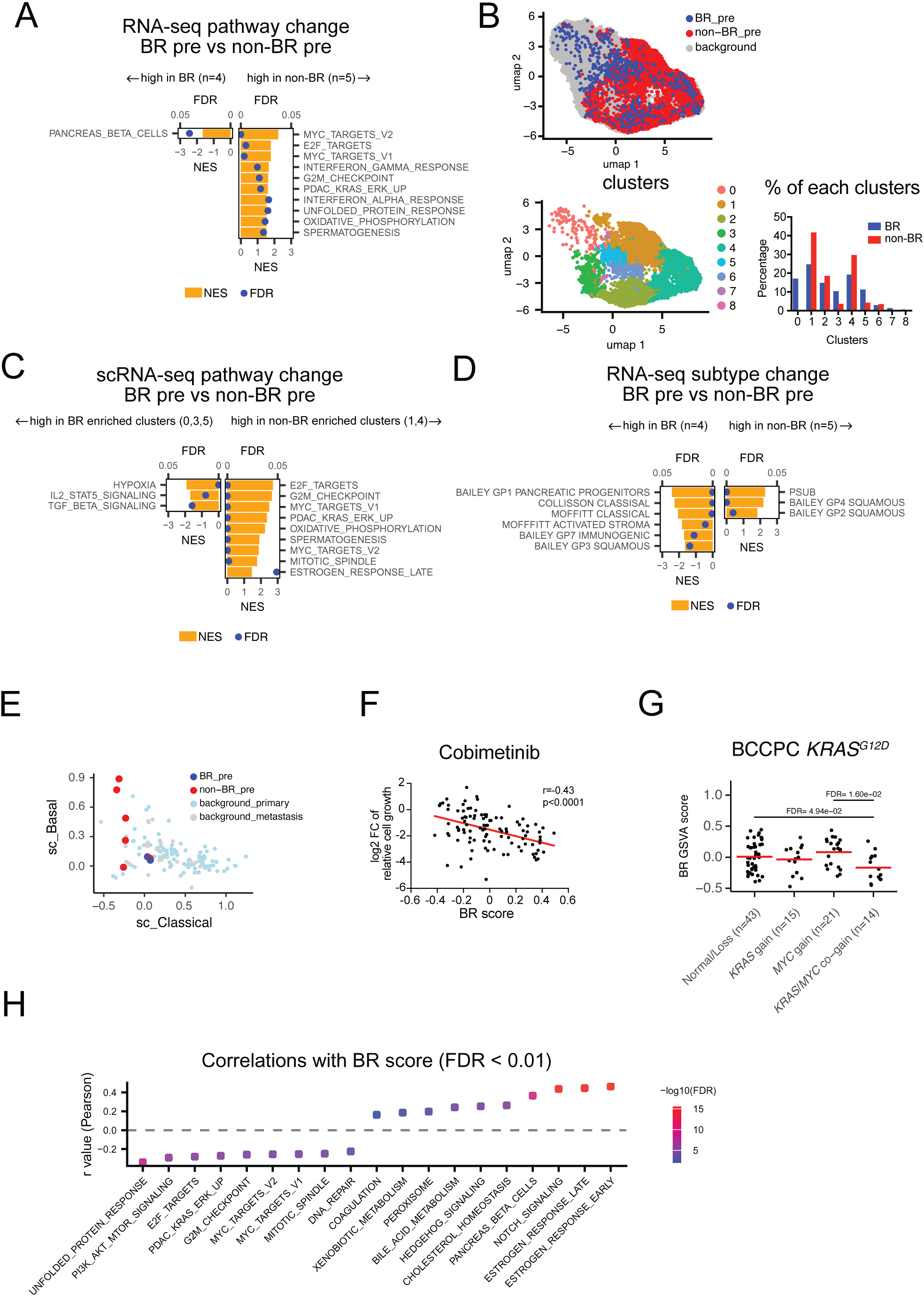
Pathways related to BR tumors and BR signature predictive of cobimetinib sensitivity. (**A**) RNA-seq pathway differences between pretreatment BR (n=4) and non-BR (n=5) tumors. GSEA of hallmark and PDAC-KRAS signatures. (**B**) (*Top*):UMAP of pre-treatment cancer cell fraction from scRNA-seq, colored by BR status (n=9). An atlas level scRNA-seq dataset was used for background mapping (Loveless IM, *et al.* 2024). (*Bottom, Left*): Unbiased clustering of cancer cell fractions. (*Bottom, Right*): Bar graph showing the percentage of each cluster in BR (n=2) or non-BR (n=7) cancer cells. (**C**) (**C**) scRNA-seq pathway differences between BR-enriched clusters (clusters 0, 3 and 5) and non-BR-enriched clusters (clusters 1 and 4). GSEA pre-rank of hallmark and PDAC-KRAS signatures. The ranking metric was defined as Log10(p value)*Log2FoldChange from MAST DEG analysis. (**D**) RNA-seq subtype differences in pre-treatment BR (n=4) and non-BR (n=5) tumors. GSEA of hallmark and PDAC-KRAS signatures. (**E**) scRNA-seq scatter plot showing subtype differences in cancer cells between BR (n=2) and non-BR (n=7). Sample level mean AddModuleScore values are plotted. Atlas level background is shown (light blue: metastasis, gray: primary) (**F**) Correlation of BR GSVA score and *in vitro c*obimetinib response across *KRAS* mutated cell lines (n=103). DepMap PRISM primary screen data. Pearson correlation. (**G**) Correlation between BR GSVA score and *KRAS*/*MYC* copy-number gain status in *KRAS^G12D^* patients. BCCPC dataset (n=93 with CNV data. *KRAS^G12D^* only, *KRAS* LOH excluded). (**H**) Correlations between the BR signature and Hallmark+KRAS/ERK pathways. BCCPC dataset (n=312 with RNA-seq data). Pearson Correlation.

### A biomarker response (BR) gene signatures predicts sensitivity to cobimetinib

To develop a gene signature to distinguish pre-treatment BR from non-BR tumors, we used RNA-seq data to identify differentially expressed genes (DEGs) and genes upregulated in BR were used as a BR signature (297 genes, log2 FoldChange >1, p <0.05, Figure S4D and Table S4). scRNA-seq showed that the BR signature was enriched in cancer cells (Figure S4E). Analysis of the independent DepMap dataset of *KRAS* mutated cell lines validated a negative correlation between BR GSVA score and resistance to cobimetinib *in vitro* (n=103, R=-0.43, p<0.0001) (Figure 3F). The BR signature did not correlate with response to other standard of care drugs including the EGFR inhibitor erlotinib, suggesting that the BR signature specifically predicts MEK inhibitor response (Figure S4F). The BR signature was significantly decreased in *KRAS^G12D^*/*MYC* co-gain tumors but was not significantly decreased in all *KRAS/MYC* co-gain tumors in the BCCPC dataset, further supporting the relationship between non-BR and *KRAS^G12D^*/*MYC* co-gain seen in WOOM samples (Figures 3G, 2J and S4G). Furthermore, the BR signature was inversely correlated with hallmark pathways including unfolded protein response, MYC targets, RS-related, DNA repair, KRAS/ERK, and PI3K/AKT (Figure 3H), suggesting these pathways contribute to MEK inhibition resistance.

### Cobimetinib biomarker response tumors have deeper KRAS and MYC pathway inhibition on treatment

To examine patient tumor molecular responses to cobimetinib, we compared post-treatment and pre-treatment paired samples. First, we compared overall post versus pre changes using bulk RNA-seq data from all paired samples. GSEA showed downregulation of KRAS/ERK, MYC, and RS-related pathways and Basal subtypes, and upregulation of metabolic pathways (Figure S5A, n=6). To focus on cancer cell-intrinsic signaling changes, we analyzed the cancer cell fraction of scRNA-seq on matched pre- and post-treatment biopsies. Using GSEA on DEGs of pseudobulk counts, comparing all paired samples post versus pre, we similarly identified downregulation of KRAS/ERK pathway, E2F target and G2M checkpoint pathways as well as upregulation of Interferon-gamma and Allograft rejection pathways (Figure S5B, n=7). Additionally, we found upregulation of the HLA class I related pathway and genes after cobimetinib treatment (Figure S5C). Subtype GSEA showed downregulation of both Basal and Classical subtype signatures in scRNA-seq with analysis of all paired samples.

Next, we focused on the difference between BR and non-BR paired post-versus pre-treatment samples. Bulk RNA-seq analysis showed downregulation of KRAS/ERK signaling, MYC targets, G2M checkpoint and E2F targets, and metabolic pathway in BR tumors compared to non-BR tumors following treatment (Figure 4A, BR:n=3, non-BR:n=3). The Bailey GP squamous signature (associated with MYC activity) was also more deeply downregulated in post-treatment BR biopsies. scRNA-seq analysis similarly identified KRAS, MYC, and RS-related pathways as significantly downregulated in tumor cells from BR compared to non-BR treated tumors (Figures 4B and C, BR:n=2, non-BR:n=5). Thus, while KRAS/ERK, MYC, E2F target and G2/M checkpoint pathways are generally downregulated with treatment (Figures S5A and S5B), they are significantly more diminished in BR compared to non-BR post- vs pre-treatment. Interestingly, scRNA-seq GSEA also showed that the intermediate basal-classical co-expressor (IC) signature^32^ was significantly downregulated in BR compared to non-BR following treatment (Figure 4C), which is consistent with an increase in the IC signature in non-BR post tumors (Figure 4D). Given that the IC signature is increased in post-neoadjuvant therapy compared with chemo-naïve patients^33^, acquisition of the IC phenotype might contribute to cobimetinib resistance.

**Figure 4:**
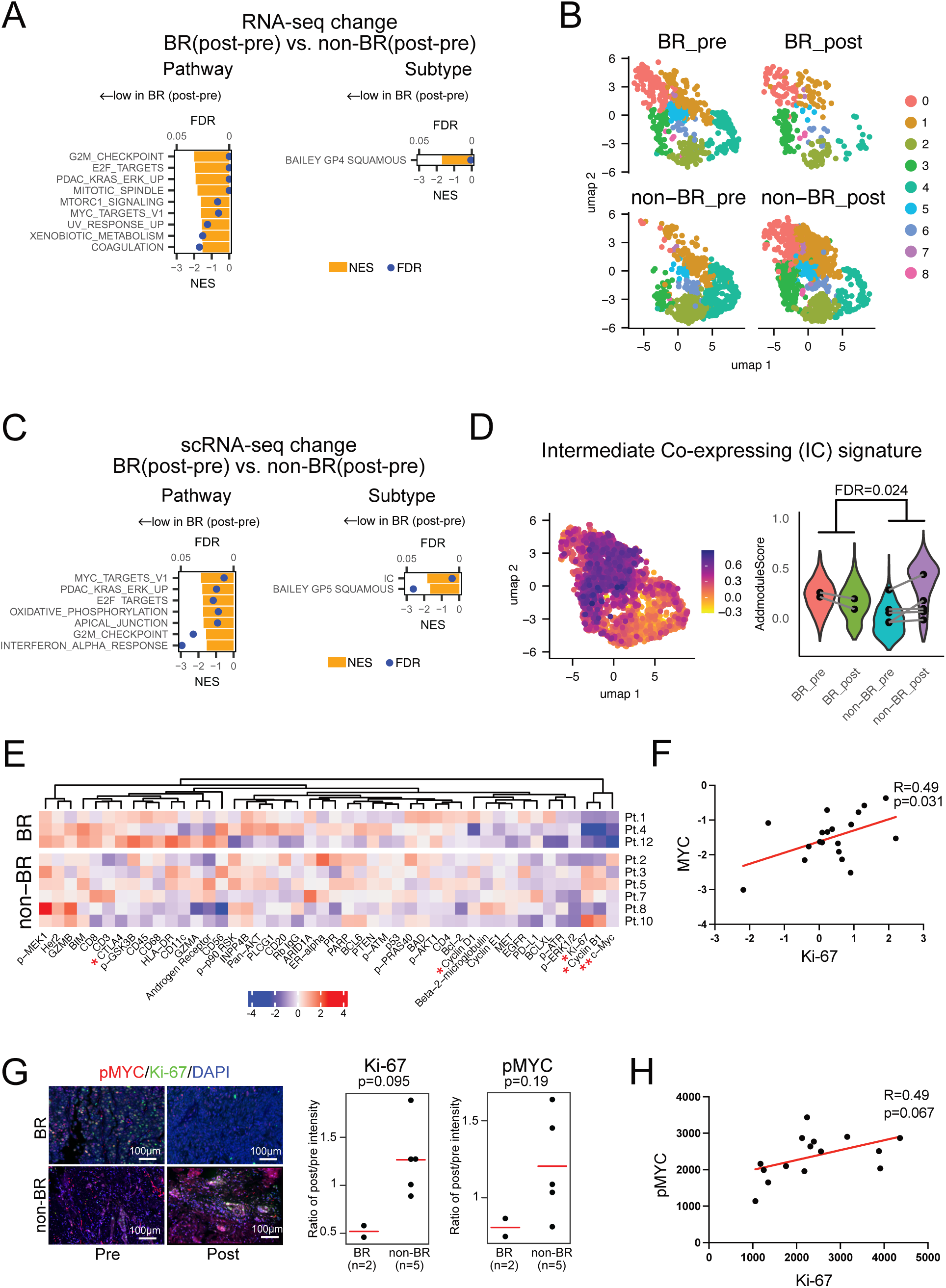
BR tumors have unique pathway changes including decrease of MYC. (**A**) RNA-seq analysis. GSEA prerank of Differentially expressed genes (DEGs) compering the post-pre difference in BR (n=2) versus non-BR (n=4) paired samples. DEGs were calculated by DESeq2. Ranking metric: log10(p-value)*Log2 fold change. (**B**) UMAP of scRNA-seq cancer cell fractions from paired pre-post samples. Plots were divided by treatment status and response status. (**C**) GSEA prerank of pseudobulk scRNA-seq DEGs comparing post-pre difference of BR (n=2) versus post-pre difference of non-BR (n=5) paired samples. DEGs were calculated by DESeq2. Ranking metric: log10(p-value)*Log2 fold change. (**D**) FeaturePlot and Violin plot of the Intermediate Co-expressor signature. Comparison of post-pre difference for BR (n=2) and non-BR (n=5). Sample level comparison of AddModuleScore. Violin plots show the cell level distribution and dots represent the sample-level mean scores. (**E**) Heatmap of post-pre changes in degital spatial profiling (DSP) normalized signal abundance in tumor cell regions. Comparing paired samples from BR (n=3) and non-BR (n=6). Wilcoxon test. *: p<0.05, **:p<0.01 (**F**) Scatterplot of DSP normalized signal abundance of KI67 and MYC. (n=19; 9 paired samples and 1 unpaired sample.) (**G**) (*Left*) Cyclic IF image showing Ki-67 and phospho-Ser62 MYC (pMYC). (*Right*): Ratio of post/pre signal intensity in tumor cells, comparing between BR (n=2) and non-BR (n=5). Wilcoxon test. (**H**) Scatterplot of Ki-67 and pMYC intensity in cyclic IF. (n=15; 6 paired samples and 3 unpaired samples stained by both markers).

DSP analysis and cycIF staining and analysis were performed to address the protein-level changes associated with treatment in BR and non-BR tumors. DSP of tumor regions revealed a significant decrease of total c-MYC protein abundance in BR tumors compared to non-BR tumors post- vs. pre-treatment, accompanied by reductions in Ki-67, Cyclin B1, and Cyclin D1 levels (Figure 4E). c-MYC was clustered together with proliferation markers Ki-67 and Cyclin B1, and expressions of c-MYC and Ki-67 were correlated between specimens (R=0.49, p=0.031) (Figure 4F). Thus, while ERK is a known regulator of MYC phosphorylation and protein stability^5,16–18^, and we observed decreased pERK throughout tumor regions and stromal regions with treatment across all paired samples (Figures 2C and S3A), and a decrease of MYC in all paired stromal regions (Figure S5D), the decrease in MYC protein with treatment in tumor regions was primarily observed in BR tumors. This suggests a discordance between inhibiting the MEK-ERK pathway and a decrease of MYC that could be a mechanism of tumor BR versus non-BR status. cycIF was used to further validate this finding. We observed decreased pSer62-MYC and concomitant reduction in Ki-67 expression in BR tumors relative to non-BR tumors; although sample level statistics with n=2 in the BR group with paired cycIF data did not reach significance (Figure 4G). pMYC and Ki-67 also tended to be correlated across all cycIF analyzed biopsies (Figure 4H). Together, this data builds on prior studies showing that pERK regulates MYC activity and protein stability via serine 62 phosphorylation^5,16,34,35^, and provides the first evidence that MYC serine 62 phosphorylation is affected by treatment with a MEK inhibitor in patients with PDAC, and highlights the potential role of bypass MYC activation in cobimetinib resistance.

### Biomarker response is associated with increased anti-tumor immunity

Next, we investigated the impact of cobimetinib treatment on the tumor microenvironment and compared the landscape of BR versus non-BR tumors. As there were few cancer-associated fibroblasts in the WOO metastatic samples examined by scRNA-seq, similar to a previous study in metastatic PDAC^31^ (Figures S6A and S6B), we focused our microenvironment analysis on immune cells (Figures 5A, S6C1 and S6D). In paired BR tumor biopsies, cobimetinib treatment increased the percentage of CD8^+^ T cells and CD4⁺ T helper (Th) cells and markedly decreased the percentage of macrophages and monocytes (Figure 5B). The increase in CD8⁺ T cells was further confirmed at the protein level by DSP analysis of the tumor-adjacent stromal region (Figure 5C). An increase of CTLA-4 in DSP tumor regions of BR patients, along with a trend toward elevated CD3 and CD8 levels might also suggest infiltration of CTLA-4 positive T cells in BR samples (Figure 4F). To further elucidate the immune cell landscape in a spatially resolved manner, we assayed the biopsies with mIHC (Figure S6E and Table S5). The mIHC stained cohort included pre- and/or post-treatment liver metastasis biopsies collected from 11 patients. The predominant immune cell populations across biopsies included T cells, antigen-presenting cells (APCs), CD11b^+^ myeloid cells, and CD163^+^ macrophages, although no significant differences in immune cell type densities were detected between BR biopsies versus non-BR biopsies (Figures S6F and S6G). Of the six pre and post paired tumors in our mIHC dataset, only one was a BR tumor, which had a distinct increase in the density of CD8^+^ T cells, CD4^+^ T helper (Th) cells, and APC cells, and a decrease in CD11b^+^ myeloid cells following treatment (Figure 5D). Although this sample pair was not analyzed by scRNA-seq, the marked immune change in this BR tumor followed similar trends to those found by scRNA-seq and DSP for other BR tumor pairs. Collectively, these findings suggest that tumors responsive to cobimetinib exhibit a remodeling of the immune microenvironment, characterized by a reduction in immunosuppressive myeloid populations and enhanced T cell infiltration.

**Figure 5:**
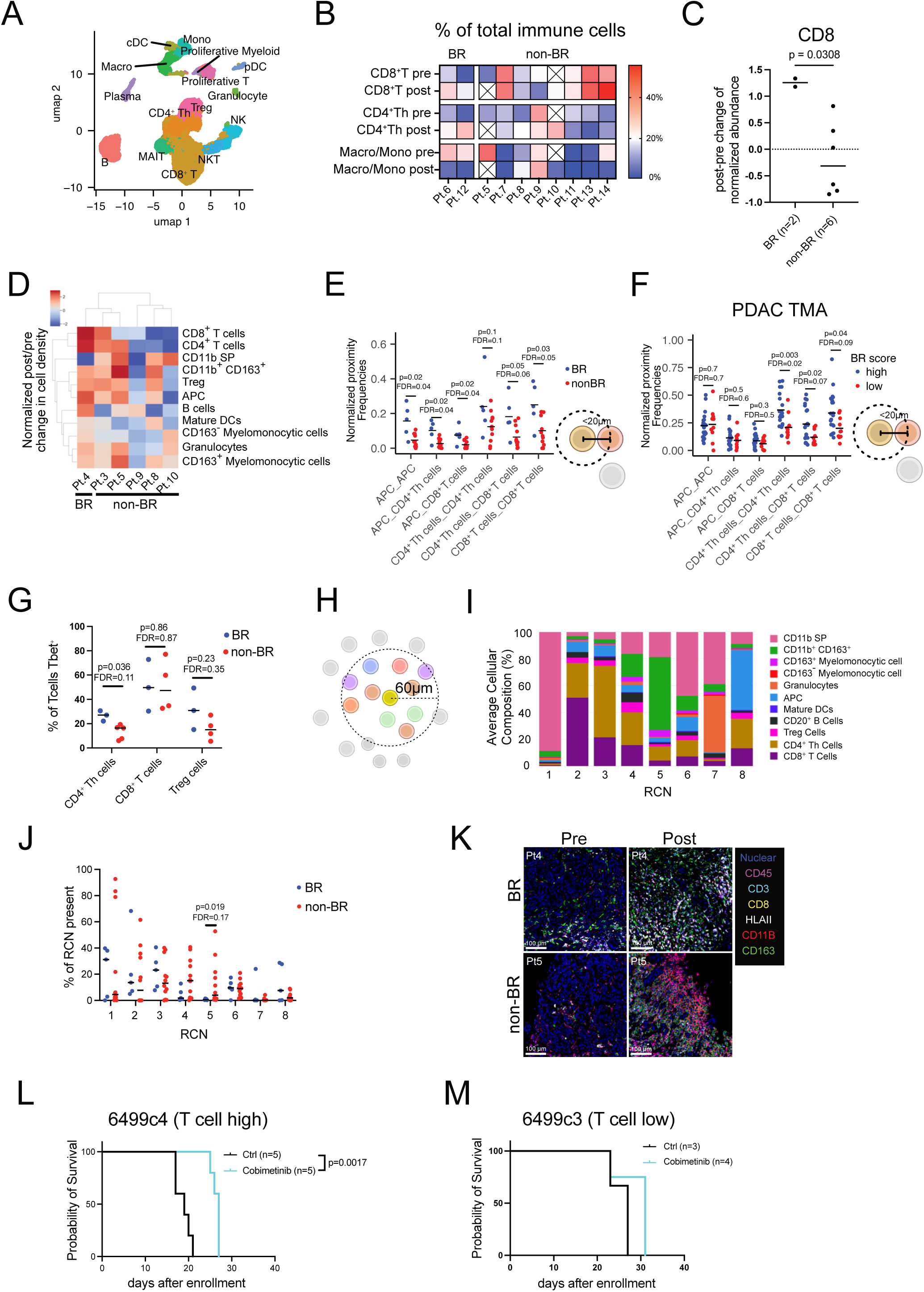
BR tumors have T cell infiltration with treatment and high T cell-APC interactions. (**A**) UMAP of WOOM scRNA-seq immune cell population colored by cell type (n=10). (**B**) Heatmap of pre- and post-treatment proportions of CD8^+^ T cell, CD4^+^ Th cell and Macrophage/Monocyte in WOOM scRNA-seq immune compartment (BR:n=2, non-BR:n=8). (**C**) DSP heatmap and CD8 signaling intensity in the tumor-adjacent stroma (BR:n=2, nonBR: n=6). Wilcoxon test. (**D**) Heatmap of the normalized density difference (post - pre) for each cell type in paired WOOM multiplex immunohistochemistry (mIHC) samples. (**E**) Cell proximity analysis (≤20μm) of WOOM mIHC. Dot plot shows proximities among Antigen presentation cells (APC), CD4^+^ Th, and CD8^+^ T cells in between BR (n=5) and non-BR (n=12). Proximity scores were normalized by cell density. Each dot represents sample-level mean across ROIs. Wilcoxon test. (**F**) Proximity analysis of APC, CD4^+^ Th and CD8^+^ T cell in PDAC TMA mIHC (n=31 patients, 20 primary PDAC, 5 liver metastases, 3 lung metastases and 3 other metastases) comparing BR high and low tumors. Proximity scores were normalized by cell density. Each dot represents sample-level mean across ROIs. Cutoff= −0.66 (details in methods). (**G**) Fraction of T-bet positive cells within each T cell subset in BR (n=3) and non-BR (n=8) WOOM mIHC samples. Each dot represents sample-level means across ROIs. Wilcoxon test. (**H**) Scheme of neighborhood analysis (≤60μm). (**I**) Neighborhood analysis (≤60μm) identifying eight immune neighborhoods in WOOM mIHC, (**J**) Fraction of each immune neighborhood in BR (n=5) and non-BR (n=12) WOOM mIHC samples. Each dot represents sample-level mean across ROIs. Wilcoxon test. (**K**) Representative pseudocolor images illustrating immune cell infiltration in BR and non-BR pre/post samples. WOOM mIHC. (**L, M**) Survival plot of cobimetinib treatment in orthotopically injected PDAC models of 6499c4 T cell high KPC cell line (**L**) and 6499c3 T cell low KPC cell line (**M**).

Based on the predominant increase in T cell and APC cell densities seen in BR tumors with treatment, we further investigated whether the spatial organization of these cell types was also related to BR status. To quantify spatial organization, we calculated the number of times CD8^+^ T cells, CD4^+^ Th cells, and APCs were within 20 µm from one another in a pairwise manner and then normalized these counts by the densities of the cells involved in the pairwise proximity^36,37^. We found the frequencies of spatial proximities involving APC-APC, APC-CD8^+^ T, APC-CD4^+^ Th, and CD8^+^ T-CD8^+^ T cell pairs were significantly higher in biopsies from BR tumors compared to non-BR tumors, regardless of pre- or post-treatment status (Figure 5E). CD8^+^ T cells undergo activation via direct contact with APCs and CD4^+^ Th cells^38^, and thus our spatial proximity results indicate the potential presence of T cell activation in BR tumors. To validate this finding, we ran a similar spatial proximity analysis on our independent PDAC TMA^3^ with mIHC analysis (n=31) for which we also had RNAseq from the same FFPE blocks, enabling us to calculate BR transcriptional scores and bin samples into BR score high versus low groups for these samples. Proximities of CD4^+^ Th-CD4^+^ Th, CD4^+^ Th-CD8^+^ T, and CD8^+^ T-CD8^+^ T cells were significantly enriched in samples with high BR transcriptional scores, although APC related proximities did not show this (Figure 5F). Notably, our TMA dataset included primary PDACs (n=20), as well as metastases from liver (n=5), lung (n=3), and other sites (n=3), indicating that T cell proximities may occur in favorable tumor microenvironments associated with a high BR score in both primary and metastatic settings. Investigation of proximities split into primary PDACs and liver metastases trended toward significance for T cell proximities in both primary and liver metastases and for APC proximities in liver metastases despite low sample numbers, as seen in our WOO mIHC (Figure S6H). Finally, we observed that BR score significantly positively correlated across all samples in the TMA with CD8^+^ T-CD8^+^ T and CD4^+^ Th-CD8^+^ T cell proximities and trended with CD4^+^ Th-CD4^+^ Th cell proximities (Figure S6I).

To further examine T cell function, we conducted a follow-up experiment and stained a subset of our WOOM samples (n=8) with T-bet, which is a transcription factor associated with type 1 immune responses, including effector T cell functions^39^. We found that BR tumors had higher fractions of CD4^+^ Th cells expressing T-bet than non-BR tumors, while T-bet expression in CD8^+^ T cells and Tregs was not significantly different between BR and non-BR tumors (Figure 5G). Collectively, our results indicate that BR tumors show evidence of T cell activation and effector function, which may be driven in part by the increased presence of CD4^+^ Th1 effector cells in BR tumors compared to non-BR tumors, further supporting the potential better biology of these tumors.

Finally, to broaden our spatial analysis to incorporate all immune cell phenotypes, we performed a recurrent cellular neighborhood (RCN) analysis, which identifies compositionally distinct spatial groupings of cells within a 60μm radius recurring across samples^37^. Our analysis resulted in eight RCNs, each composed of different immune cell phenotypes (Figures 5H and 5I). Notably, RCN 5, characterized by the presence of immunosuppressive CD163^+^ macrophages, was present more often in non-BR tumors, compared to BR tumors (Figures 5J and S6J). Overall, our spatial results indicate that both increased lymphoid and decreased myeloid cell spatial architectures contribute to establishing a favorable tumor immune microenvironment in the BR tumor cohort (Figure 5K).

To test the hypothesis that T-cell infiltration is a key determinant of cobimetinib response, we orthotopically implanted syngeneic single cell-cloned mouse PDAC cell lines characterized with high or low T-cell infiltration ability^40^, derived from the same tumor, into the pancreatic tail of syngeneic host mice and compared their responses to cobimetinib treatment (Figures. 5L and M). We found that T cell high tumors exhibited a greater response to cobimetinib treatment compared to T cell low tumors, supporting a role for T-cell infiltration in cobimetinib efficacy.

### Correlation between cobimetinib and KRAS inhibitor responses in models of BR and non-BR tumors

Since their initial discovery, several KRAS^G12D^ mutant-specific inhibitors have undergone clinical assessment, including MRTX1133^41^. Given that non-BR PDX displayed resistance to MRTX1133 (Figure 2H), we sought to investigate the concordance of MEK-inhibitor and KRAS-inhibitor resistance. Analysis of the DepMap database revealed a correlation between cobimetinib and MRTX1133 responses across 32 *KRAS^G12D^* mutant cell lines, irrespective of cancer type (r = 0.46 p = 0.0077; Figure 6A). Moreover, our BR signature anti-correlated with reduced sensitivity to MRTX1133 in the DepMap dataset (r = −0.49, p= 0.0055; Figure 6B). Furthermore, *KRAS*/*MYC* co-gain cell lines were less sensitive to MRTX1133 (p=0.0478; Figure 6C). A correlation was also found between cobimetinib and the pan-KRAS inhibitor RMC7977 response (r = 0.501, p=1.29e-07; Figure S7A) and between BR score and RMC7977 response (r = −0.387, p=7.42e-05; Figure S7B) across all *KRAS* mutated cell lines. To validate these findings in our low passage patient-derived cell lines (PDCLs) generated from the WOO pre-treatment biopsies, we treated four *KRAS^G12D^* mutated PDCLs with MRTX1133 and found that the two PDCLs with *KRAS*/*MYC* co-gain were most resistant (Figure S7C).

**Figure 6:**
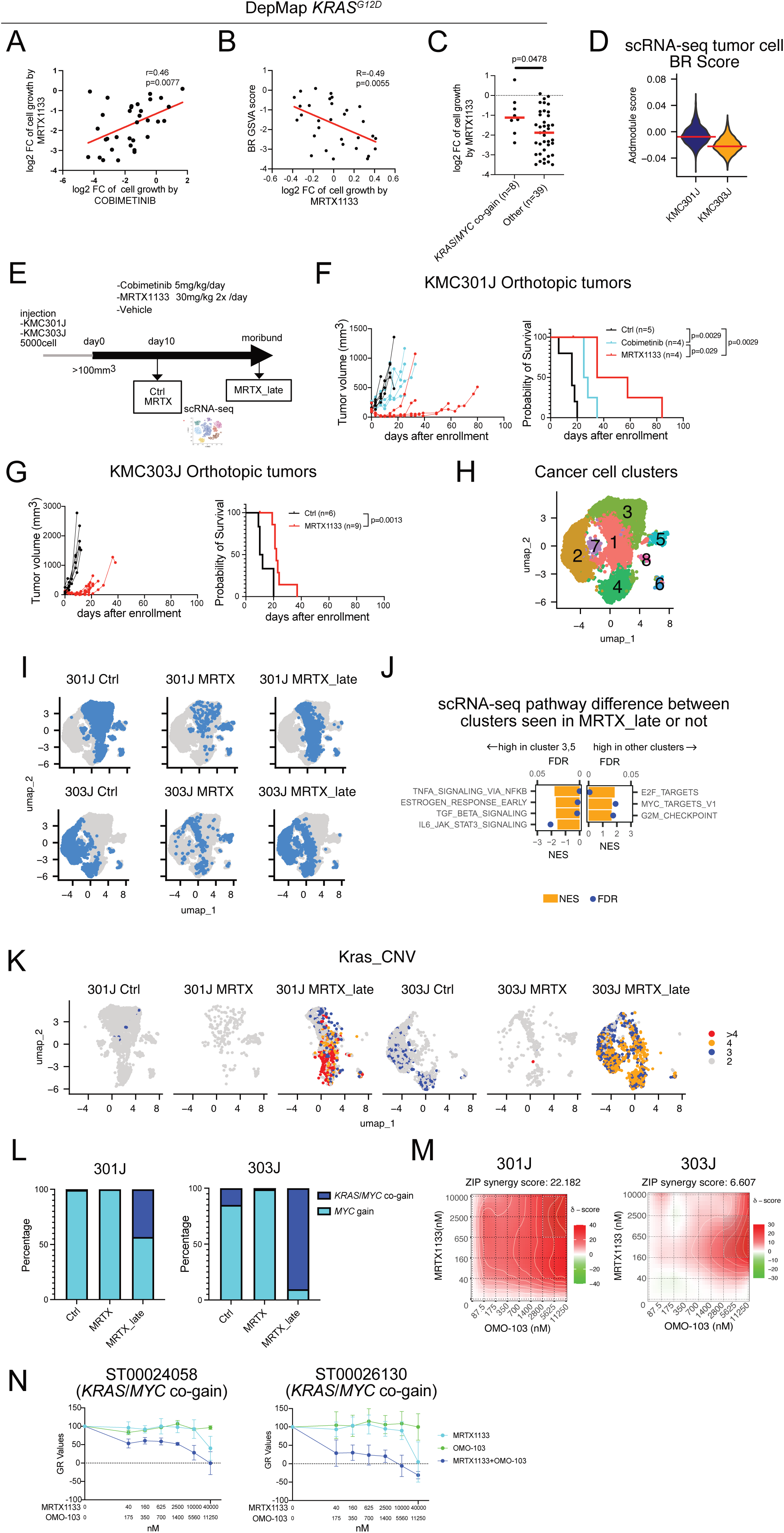
Models of BR and non-BR tumors demonstrate mechanism of KRAS inhibitor resistance. (**A**) Correlation between cobimetinib response and MRTX1133 response in *KRAS^G12D^* mutated cell lines (n=33). DepMap. (**B**) Correlation between MRTX1133 response and BR signature GSVA score in *KRAS^G12D^* mutated cell lines (n=46). DepMap. (**C**) Dot plot of MRTX1133 response by relative CNV status. *KRAS*/*MYC* co-gain (n=8) and other CNV categories (n=39). DepMap. (**D**) BR signature score in tumor cell from 301J and 303J orthotopic tumors. scRNA-seq AddmoduleScore. Cell-level Wilcoxon test. (**E**) Schematic of the *in vivo* drug treatment study. Samples for scRNA-seq were collected after 10 days of treatment (Ctrl, MRTX) or at moribund state (MRTX_late). (**F**) Tumor growth curve and survival analysis of cobimetinib and MRTX1133 treatment in KMC301J orthotopic tumors. (**G**) Tumor growth curve and survival analysis of MRTX1133 treatment in KMC303J orthotopic tumors. (**H,I**) UMAP of tumor cell fractions from scRNA-seq. (**H**) UMAP clustering. (**I**) UMAP splitted by treatment conditions for each cell lines. (**J**) GSEA pre-rank of hallmark and PDAC-KRAS/ERK signature comparing tumor cell clusters present versus not present in MRTX_ late. Clusters 3 and 5 versus other clusters. Rank metric; Log10(p-value)*-log2(fold change) from of MAST DEG analysis. (**K**) FeaturePlot of *Kras* CNV number infereed from infer CNV. scRNA-seq. (**L**) *KRAS*/*MYC* CNV status in each tumor across treatment conditions. (Left: 301J Right: 303J). (**M**) Combination treatment of MRTX1133 and OMO-103 in KMC301J and KMC303J cell lines. Percent inhibition was calculated by GR metrics and Synergy score computed using ZIP method. (Left: 301J Right: 303J). (**N**) Combination treatment with MRTX1133 and OMO-103 to *KRAS*/*MYC* co-gain patient derived cell lines. Growth rate inhibition metrics (GR metrics) was used.

To investigate BR and non-BR tumors in an *in vivo* laboratory model, we took advantage of our KMC^ERT^^2^ mouse model of adult-onset acinar cell-derived PDAC, driven by coordinated *Kras^G12D^*and deregulated *MYC* expressed from the *Rosa26* locus (*Ptf1aCreER; Kras^G12D^; Rosa26^MycWT/MycWT^*), representing heterogeneity of human PDAC subtypes leading to metastatic PDAC^42^. Analysis of scRNA-seq data from two KMC^ERT^^2^ orthotopic tumors (KMC301J and KMC303J) exhibited differential BR signature scores (Figure 6D). We assessed response to cobimetinib and/or MRTX1133 in the KMC301J and KMC303J syngeneic models and analyzed by scRNA-seq (Figure 6E). Cobimetinib or MRTX1133 prolonged survival, but only MRTX1133 induced regression of KMC301J tumors. Notably, while all the tumors initially regressed with MRTX1133 treatment, they eventually relapsed, even on treatment. There were inter-individual differences in time to recurrence, reflecting heterogeneity in the model and differential capacity for acquired resistance (Figure 6F). For KMC303J tumors, we decided to treat only with MRTX1133 based on their lower BR signature and the reduced response to cobimetinib compared to MRTX1133 in the KMC301J tumors. While KMC303J tumors initially responded to MRTX1133, they relapsed within 10–15 days, suggesting some degree of innate resistance to MRTX1133 within this model (Figure 6G). All together supporting the connection between BR score and sensitivity to KRAS inhibition both *in vitro* and *in vivo*.

### Models of BR and non-BR show similar resistance mechanism to WOO patient tumors

Analysis of the tumor cells from scRNA-seq of the MRTX1133-treated KMC301J and KMC303J tumors (Figures S7D and 6H) showed that untreated KMC301J and KMC303J tumor cells were distinct, but after MRTX1133 treatment, they converged into some overlapping clusters that persisted through to termination (Figure 6I). Interestingly, KMC301J tumor cells changed some of their cluster populations between non-treatment and relapsed tumors, while KMC303J tumor cells mostly populated similar clusters on relapse. This further supports KMC301J and KMC303J as models of acquired resistance and innate resistance, respectively. At relapse, tumor clusters 3 and 5 diminished in 301J tumor cells, and these populations were not present in KMC303J tumor cells (Figure 6I), prompting a comparative analysis of these sensitive clusters with other resistant clusters. Pathway analysis showed enrichment of NFκB, TGFβ and JAK/STAT pathways in the sensitive clusters and MYC and RS-related pathway upregulation in resistant tumor clusters (Figure 6J), similar to non-BR patient-derived tumors from the WOOM trial (Figure 3C). Infer CNV analysis revealed that *Kras* gain on chromosome 6 was increased at relapse in both KMC301J and KMC303J tumors (Figure 6K). Given the KMC^ERT^^2^ cell lines have 2 copies of external *Rosa^Myc^*alleles by design, this *Kras* gain acquisition is consistent with a *Kras*/*Myc* co-gain status in tumor relapse (Figure 6L). This suggests *Kras*/*Myc* co-gain is also a potential resistant mechanism in our KMC^ERT^^2^ PDAC mouse model. Since oncogenic RAS and MYC inhibit expression of MHC class I in tumor cells^43,44^, we investigated the activity of MRTX1133 in modulating antigen presentation in KMC301J and KMC303J tumor cells. After 10-days of MRTX1133 treatment, KMC301J tumors, but not KMC303J tumors, exhibited an increase in MHC class I gene, H2-K1 (Figure S7E). In contrast, the DC cell population and MHC class II expression in myeloid cells was increased in both tumors (Figure S7F). Lymphoid cell infiltration, however, was observed only in KMC301J tumors, suggesting that tumor-intrinsic MHC class I activation may contribute to lymphoid cell recruitment in the KMC301J BR tumor model (Figure S7G).

### Concurrent KRAS and MYC inhibition overcomes therapeutic resistance

From the finding that *KRAS*/*MYC* co-gain is a resistance mechanism to targeting the KRAS/MEK pathway, we next examined the combination of MRTX1133 with the MYC inhibitor, OMO-103, that is being assessed in clinical trials^45^. Indeed, the combined inhibition of MRTX1133 and OMO-103 had a synergistic effect *in vitro* in both KMC301J and KMC303J models (Figure 6M). To assess if this combination could overcome the therapeutic resistance mediated by *KRAS*/*MYC* co-gain, we next treated two PDCLs from the WOO trial that harbor a *KRAS^G12D^* mutation and *KRAS*/*MYC* co-gain with MRTX1133 and OMO-103. Even using sub-therapeutic dosing of OMO-103, we found that together with MRTX1133, the combination demonstrated synergistic activity in both cell lines (Figure 6N). These laboratory models strongly support: 1) our BR signature as a marker of KRAS inhibitor sensitivity, 2) *KRAS*/*MYC* co-gain in PDAC as a resistance state, and 3) the potential for concurrent KRAS and MYC inhibition to overcome resistance to therapeutic targeting of the KRAS/MEK pathway.

## Discussion

In this study we identified the prevalence of *KRAS* and *MYC* copy number co-gains in PDAC. While single gains of either *KRAS* or *MYC* were not predictive of survival, a *KRAS/MYC* co-gain was an independent predictor of worsened survival in patients with PDAC. Notably, the frequency of *KRAS*/*MYC* co-gain increased in the metastatic setting, suggesting a role in PDAC progression. Moreover, our data extends the emerging understanding that distinct *KRAS* mutant allele number and mutational variance (e.g., *KRAS^G12D^* vs *KRAS^G12V^*or *KRAS^Q61H^*) correlates with differential patient survival outcomes^14^. Mechanistically, we found that, unlike *KRAS^G12V^* and *KRAS^G12R^,* copy number gain of *KRAS^G12^*^D^ mutation is associated with activation of the KRAS/ERK and MYC pathways, and PI3K/MTORC signaling. Our data further elucidates the importance of *MYC* gain in the activation of these pathways. Together, our findings highlight the significance of *KRAS*/*MYC* co-gain in driving metastatic PDAC while underscoring the mutation variant-specific roles of *KRAS* gain as well as the function of *MYC* gain in the feedforward activation of KRAS/ERK and PI3K tumor-promoting pathways.

Using our WOO trial platform, which integrates multi-omics analyses and patient-derived models to assess biomarker-defined responses, we interrogated the biological activity of the MEK inhibitor, cobimetinib, in patients with metastatic PDAC. Our results indicate that a lack in tumor-responsiveness to cobimetinib (non-BR) was characterized by high activation of the KRAS/ERK and MYC pathways, with enrichment for *KRAS* and *MYC* gene gain and co-gain in *KRAS^G12D^* mutated tumors. Furthermore, by applying a biomarker response (BR) transcriptomic signature generated from the pre-treatment tumors of this WOOM trial to our independent PDAC dataset we identified a strong correlation between low BR signature and activation of the KRAS/ERK, MYC and PI3K pathways, and *KRAS^G12D^*/*MYC* co-gain status. Together, this supports the contention that *KRAS^G12D^*/*MYC* co-gain–mediated cooperative activation of the KRAS and MYC oncogenic pathways contributes to tumor resistance to cobimetinib.

A key strength of the WOO trial is its ability to rapidly elucidate drug-induced tumor-intrinsic and microenvironmental changes in patients. By comparing pre- to post-treatment tumor response, we demonstrated that MYC pathway downregulation, including pMYC depletion, correlates with biomarker response. While the regulation of pMYC by pERK is well established, this study provides the first evidence of downstream uncoupling of these pathways as a resistance mechanism in PDAC tumors in humans. Interestingly, basal subtype signatures were suppressed irrespective of biomarker response status with treatment, suggesting that signaling through the MEK/ERK pathway, also universally suppressed, rather than MYC, plays a dominant role in maintaining basal transcriptional networks. In contrast, the Intermediate basal-classical co-expressor signature was distinctively upregulated after treatment in non-BR tumors. This suggests that acquisition of an intermediate subtype phenotype contributes to tumor resistance to cobimetinib treatment.

Our study also highlights the interplay between BR status and the tumor immune microenvironment. We observed treatment-induced MHC class I activation of the tumor cells with CD4^+^ and CD8^+^ T-cell infiltration in the BR tumors and this increase in MHC class I and lymphoid cell infiltration was recapitulated in 10 days KRAS inhibitor treated KMC mouse model of BR tumor. Additionally, BR tumors exhibited a more favorable immune microenvironment, characterized by enriched APC, CD8^+^ T, and CD4^+^ Th cell spatial interactions and a reduced immunosuppressive CD163^+^ macrophage enriched neighborhood. The association between predicted BR status by transcriptomic signature and T cell-T cell spatial interactions was validated in an independent PDAC mIHC dataset. The importance of T cell infiltration in response to MEK inhibition was also shown by *in vivo* mouse models of PDAC which display different T cell infiltration. Since, the presence of APC, CD8^+^ T, and CD4^+^ T-cell spatial proximity is known to predict response to immune checkpoint blockade (ICB) therapy^38,46^, enrichment in this triad formation in BR tumors would support future clinical trials combining RAS/ERK/MYC pathway inhibitors with ICB, which has also been suggested in mouse studies^47,48^.

The emergence of KRAS inhibitors as a viable therapeutic strategy is poised to alter the treatment landscape for patients with PDAC^11,49–51^. Even so, resistance to these targeted agents is a likely reality that requires forethought as to the compensatory mechanisms driving resistance, and a need for identifying combination strategies that improve the response to KRAS inhibitors. To this end, our development of a BR signature for cobimetinib from the WOO trial demonstrated translation to KRAS inhibitor sensitivity in the external DepMap data and in our mouse models recapitulating BR and non-BR phenotypes. Notably, we identified enrichment of KRAS/ERK, MYC, and replication stress related E2F and G2M checkpoint pathways, as well as *Kras*/*Myc* genetic co-gain in KRAS inhibitor resistance in relapsed tumors in our genetic mouse model, suggesting that insights gained from our MEK inhibitor clinical trial could inform strategies for patient selection and approaches to overcome resistance to KRAS-targeted therapies. Moreover, these findings support the concept that concurrent targeting of KRAS and MYC could overcome this resistance mechanism.

A limitation of this study is the relatively small patient sample size and lack of long-term response assessment in the WOO trial, which may affect the characterization of response and the generalizability of our findings. To mitigate this, we leveraged PDX models developed from patients’ tumors in the trial as a proxy for long-term response and applied our novel biomarker response signature to external large-scale cobimetinib and KRAS-inhibitor treated KRAS-mutant cell line datasets. In addition, validations in independent human PDAC datasets and genetic mouse models for molecular and cellular response characteristics were conducted. We were also clinically limited to single site biopsy specimens, and thus the effect of intra-tumor heterogeneity could not be fully addressed. However, our study of clinical and laboratory responses to KRAS-MEK inhibition contributes molecular and cellular tumor and immune response data that may assist in the design of therapeutic clinical trials with multi-omics profiling to validate these findings and refine biomarker-driven therapeutic strategies for PDAC.

## Methods

### BCCPC cohort

The human PDAC dataset used in this study was extended from our recent paper^3^. From a de-identified dataset of patients diagnosed with and/or treated for PDAC at our institution between 2004 and 2025, we identified 308 patients for which we had specimens with CNV data and 312 patients for which had RNA-seq data. 287 patients were overlapped with CNV data and RNA-seq data. Patients whose primary tumor was located at the ampulla of Vater but classified as pancreatobiliary subtype were included (*n*L=L9). Clinical course time points, stage, grade, nodal involvement, resection margins and angiolymphatic invasion were provided as de-identified data by the OHSU cancer registry with quality control data verification by pathologists (B.B. and T.M.). Patient demographics were also collected and include age and self-reported sex. To adhere to our clinical definition, we did not exclude patients from any cohort due to short survival. All patient information was frozen in February 2025.

### RNA-seq and Genomic Alteration panel processing

FFPE tissue of patients’ biopsy are scraped for DNA and RNA extraction. Solid tumor total nucleic acid was extracted from these tumor regions using Chemagic 360 sample-specific extraction kits (Perkin Elmer) and digested by proteinase K. RNA was purified from the total nucleic acid by DNase-I digestion. DNA sequencing of 596 genes and whole-transcriptome RNA sequencing were performed by Tempus as previously described^3^.

Briefly, 100 nanograms (ng) of DNA for each tumor sample was mechanically sheared to an average size of 200 base pairs (bp) using a Covaris Ultrasonicator. DNA libraries were prepared using the KAPA Hyper Prep Kit, hybridized to the xT probe set, and amplified with the KAPA HiFi HotStart ReadyMix. One hundred ng of RNA for each tumor sample was heat fragmented in the presence of magnesium to an average size of 200 bp. Library preps were hybridized to the xGEN Exome Research Panel v1.0 (Integrated DNA Technologies) and target recovery was performed using Streptavidin-coated beads, followed by amplification with the KAPA HiFi Library Amplification Kit. The amplified target-captured DNA tumor library was sequenced using 2×126bp PE reads to an average unique on-target depth of 500x and 150x (normal) on an Illumina HiSeq 4000. The amplified target-captured RNA tumor library was sequenced using 2×75bp PE reads to an average of 50M reads on an Illumina HiSeq 4000 or NovaSeq6000. Samples were further assessed for uniformity with each sample required to have 95% of all targeted bp sequenced to a minimum depth of 300x.

### DNA sequence analysis

DNA Variant detection, visualization, and reporting were performed as previously described^3^. Alignment and mapping were to GRCh37 using Novo align + BWA. CNVs derived from proprietary tumor/normal match analysis using CNAtools and CONA^52^. Statistics are Fisher’s Exact test to look for significantly changed alteration rates between cohorts.

### RNA-seq data processing

Paired-end fastq sequences were trimmed using Trim Galore (v.0.6.3) and default parameters. Pseudoalignment was performed with kallisto (v.0.44.0) using genome assembly GRCh38.p5 and GENCODE (v.24) annotation; default parameters were used other than the number of threads. The Bioconda package bioconductor-tximport (v.1.12.1) was used to create gene-level counts and abundances (TPMs). Quality checks were assessed with FastQC (v.0.11.8) and MultiQC (v.1.7). Quality checks, read trimming, pseudoalignment and quantitation were performed via a reproducible snakemake pipeline, and all dependencies for these steps were deployed within the anaconda package management system. Batch correction was performed in TPM level to remove batch effect based on platform difference between older samples and newer samples. Genes unnormalized count>=10 at least one-third of total number of samples were included. DEseq2 (v.1.44.0) was used for obtain variance stabilizing transformation (VST) counts or Differentially Expressed Genes. Raw BCL files were used to generate FASTQ files using BCL2FASTQ (v2.17). Adapters were trimmed using Skewer (v0.2.2). Reads were then aligned with STAR (v.2.5.4a) to generate BAM files and undergo UMI deduplication with umitools (v1.0.1). Following deduplication, samples were converted to FASTQ files using bedtools (v2.27.1) before undergoing quantification using Kallisto (v0.44.0) with the Ensembl GRCH37 reference transcriptome. Raw RNA expression data were normalized to minimize the effect of technical artifacts such as GC content, transcript length, and library size^52,53^. Batch correction was performed to account for small technical differences between two versions of the RNA-seq assay, including a modified exome capture probe set and the addition of UMIs^54^. Genes unnormalized count>=10 at least one-third of total number of samples were included.

### External datasets

TCGA PAAD firehose dataset with CNA and RNA-seq data was obtained from cBioportal and filtered for high tumor purity PDAC samples (n=150). CPTAC PAAD GDC dataset with CNA data was obtained from cBioportal (n=95, n=85 with RNA-seq data). UTSW PAAD dataset with CNA data was also obtained from cBioportal (n=109).

### Survival analysis

Survival of the BCCPC cohort was calculated as the interval from date of sample collection to date of death or last follow-up. were excluded from survival analyses. Only patients alive 30 days or more after surgery were included to avoid analyzing death related to surgical complications. All *P* values and adjusted HRs associated with survival analyses are from multivariable Cox proportional hazards models accounting for covariates including Stage, Tumor grade, KRAS variant. Covariates were selected based on statistically significant univariate associations with OS. Analyses were conducted using graph pad prism (v.10.1).

### GSVA

Batch corrected log2(TPM+1) was used for calculating GSVA score with R library GSVA (v.1.52.3). MSigDB Hallmark gene set, PDAC-KRAS-ERK signature^15^, BR signature defined in this paper, and pORG score^3^ were used.

### Virtual Inference of Protein-activity by Enriched Regulon analysis (VIPER)

The transcriptional regulon enrichment analysis was performed using VIPER with the TCGA PAAD ARACNe-inferred network^55,56^. Batch corrected log2 (TPM+1) data were used for running VIPER. VIPER regulon scores were used for cohort comparisons.

### Summary of WOO sample preparation

Tumor and blood samples were collected from study participants in the Window of Opportunity trial (NCT04005690) under OHSU Institutional Review Board approval #00019211. Informed consent was obtained, and all protocols were performed in accordance with federal and state regulations, and policies outlined by the OHSU IRB. Tumor tissue from patients enrolled in the Window of Opportunity (WOO) clinical trial was collected from metastases during scheduled biopsy procedures, both prior to and following 10 days of treatment. Treatments were performed following a minimum 10-day washout of prior therapy. A proportion of the samples was dispatched for omic analysis and pathology, while the remainder was transferred to the laboratory for modelling. Samples were split into multiple models; a small whole viable sample is implanted as a patient-derived xenograft (PDX) into an NOD.Cg-Prkdcscid Il2rgtm1Wjl/SzJ (NSG) mouse (CAT#5557, The Jackson Laboratory,) mouse. The remaining tumor tissue underwent disaggregation using gentleMACS kits (Miltenyi Biotec, #130-095-929) in accordance with the manufacturer’s protocol. The resultant lysate was resuspended and filtered through a 70-µm cell strainer (130-098-462; Miltenyi Biotec). The cells were then collected by centrifugation (300 × g for 7 min at 4 °C) and resuspended at a concentration of 700-1200 cells/µl. The isolation of live cells was conducted using the EasySep Dead Cell Removal (Annexin V) Kit (STEMCELL Technologies, #17899), which was employed for single cell RNA-seq. The cells were subsequently cultured to establish a cell line in continuously regenerating cell (CRC) medium. In addition, a small whole viable tissue was cryopreserved in liquid nitrogen for future experimental use. Blood was collected on the days of tumor biopsy, prior to tumor biopsy collections. Blood was allowed to clot for 30 min and then processed 1500g for 10 min to isolate serum.

### CA 19-9 analysis

The OHSU Hospital Core laboratory quantified the level of CA 19-9 in serum samples using Beckman Coulter GI Monitor, CA 19.9 two-site immunoenzymatic sandwich assay.

### Immunostaining

Five-micron sections of tumor biopsies were subjected to immunohistochemistry using Ventana autostainer in a CLIA-certified histology laboratory. Ki-67 was evaluated using the VENTANA CONFIRM anti-Ki-67 (30-9) Rabbit Monoclonal Primary Antibody (Ventana Medical Systems, Inc., Tucson, AZ). All stained sections, including H&E slides, were scanned using a Leica AT2 digital scanner. Ki-67 positive percentage in tumor cells was assessed by pathologist.

### mIHC Staining and Image Analysis

Sequential IHC was performed on 5um FFPE sections using a previously published protocol^57^. Briefly, slides were deparaffinized and stained with hematoxylin followed by whole-slide scanning at 20x on an Aperio AT2 (Leica Biosystems). Tissues were antigen retrieved in pH 6.0 citrate solution, then stained and stripped sequentially with the antibodies (Table S6). Image processing was performed using previously described methods^58^. All available high-quality tissue, excluding necrosis, normal tissue, and damaged tissue were used for downstream analysis. Single marker binary thresholds were set using FCS Express Image Cytometry RUO (De Novo Software) and visually validated for positive marker expression. Single cell classification was performed using R statistical Software based on the hierarchical gating strategy in Table S5.

### mIHC Spatial Analysis

Spatial proximity frequencies and recurrent cellular neighborhoods (RCNs) were calculated following previously published methods and code using pandas (v2.2.1), numpy (v.1.26.4), scipy (v1.12.0), and scikit-learn (v1.4.2) Python software packages^37^. Briefly, proximities were calculated by counting the number of times two cells were located within 20 µm of each other for each tissue region. This number was then divided by the sum of the densities of the two cell types involved in the proximity for the given tissue region to normalize for cell types present in high abundances. Finally, all proximity frequencies were log10+1 transformed to account for proximities present in different orders of magnitude. RCNs were calculated by first defining neighborhoods for every immune cell present in the dataset by counting the immune cells within a 60 µm radius of each seed cell. Neighborhoods were then k-means clustered according to the proportions of cell phenotypes present in each neighborhood, resulting in clusters of neighborhoods—or RCNs—that consisted of similar cell types. The elbow method was used to determine the optimal number of clusters. Additionally, in the dataset, we had replicate samples of three of the biopsies, offering a unique opportunity to assess tissue heterogeneity within these biopsies. Analysis of the replicate samples showed varying degrees of heterogeneity in immune cell densities, with one of the three biopsies exhibiting large differences and the other two biopsies exhibiting smaller differences (Figure S6J), reflecting the challenge of working with small tissue areas as previously shown in primary PDAC^59^. Given that our overall goal of performing mIHC was to assess immune contexture, we included only the replicate with the highest overall immune cell density for each of the three biopsies. In the analysis of PDAC TMA, to maximize our understanding of the immune proximity differences, BR GSVA scores of corresponding RNA-seq were divided into high and low groups at each decile, and Wilcoxon tests were performed for each division, and the cutoff yielding the smallest p-value (BR score = −0.66) was defined as the optimal threshold.

### Cyclic IF

Slides with five-micron sections of FFPE tissue of biopsies were stained using the CycIF protocol ^60^. In summary, tissues were stained with 4 antibodies plus DAPI each round and were taken through six to thirteen rounds of staining (Table S6). As tissues experienced varying degrees of tissue loss, most analysis focused on the first five rounds of staining, providing information for 20 proteins. Only proteins common to all batches were retained during the data integration process across batches. After staining, image segmentation and cell identification were performed using the Cellpose algorithm. Fluorescence intensities were extracted with the MPLEXABLE pipeline and served as a proxy for protein expression. For each tissue, raw intensity values were Z-normalized on a per-protein basis; negative Z-scores were truncated to zero. Thresholds for positive marker expression were set according to channel-specific minimum signal levels, with some manual adjustments informed by visual inspection. Intensities below their respective thresholds were also zeroed. Cell-type classification relied on a weighted scoring approach. A predefined weight matrix (proteins × cell types) was multiplied by the normalized intensity matrix (cells × proteins) to generate cell-type scores. Each cell was assigned the type with the highest score. Cells with no score exceeding 0.2 were designated as “stromal” to capture unclassified or low-expression populations. The post/pre ratio of mean intensity of the cancer cells fraction was compared between the biomarker response (BR) group and the non-BR group. Pt.1 biopsies were stained by two different batches and old batch data was used for Ki-67 staining post-pre comparison, because Ki-67 staining of Pt.1 post treatment new batch was failed.

### NanoString Digital Spatial Profiling (DSP) assay

FFPE core biopsy samples were deparaffinized with Xylene, 100%, 95% and 70% ethanol sequentially. Antigen retrieval was performed with sodium citrate buffer (pH6.0) under high pressure for 15 min. After blocking with Buffer W for 1 hour, samples were incubated overnight at 4°C with NanoString’s protein modules of Immune core cell profiling, Pan-tumor, Cell death, PI3K/AKT and OHSU custom protein panel which mainly focuses on cell cycle and DNA damage signaling pathways (panel version 1.2 was used for Pt1_pre to Pt12_pre samples and panel version 2.1, which was modified from version 1.2, was used for Pt12_pre and Pt12_post samples). Pan-Cytokeratin (PanCK, AF532) and CD45 (AF594) were co-incubated for solid tumor morphological markers. SYTO13 was used for DNA staining. Tumor-enriched regions were reviewed by a board-certified pathologist and three regions of interest (ROIs) per tissue were collected for molecular profiling. Each ROI was segmented into two distinct areas of interest (AOI) corresponding to the tumor region and an adjacent immune region 20mm from the tumor. UV-cleaved DNA oligos from antibody panel were collected in 96-well plate and hybridized with 6-color fluorescent barcoded hyb codesets for nCounter read. Hybridized tagsets (DNA oligo + codesets) were processed and immobilized in the cartridge for molecule count with nCounter MAX system. nCounter data was re-imported to DSP and used to generate an OHSU clinical report. Briefly, a TMA control consisting of 15 cell lines was tested in each run, and abundance data was input into an RUV-based normalization method^61^ to reduce batch effects. For clinical specimens, abundances from tumor and peritumoral segments were analyzed separately. Normalized antibody abundances from the 3 AOI’s were averaged and plotted as percentiles relative to the expression levels of an internally established reference cohort (6 cases) of pancreatic carcinomas. For comparing two serial biopsies (pre- vs on/post-treatment) from the same patient, the signal difference in each antibody was compared to an antibody-specific threshold for assay variability. A difference was reportable when it exceeded the 95^th^ percentile of the delta distribution from a pairwise assessment of technical replicate TMA controls. Taking account of batch effect for normalization, only panel v1.2 result was used for the sample level analysis (Figures 2C, 4F, S3A and S5D), and both panel v1.2 and v2.1 results were used for the analysis of delta (post-pre) (Figures 4E and 5C).

### Gene Set Enrichment Analysis (GSEA)

Counts data of pre samples are extracted from WOOM cohort (n=4 for BR, n=5 for non-BR) and filtered to retain the genes which have more than 10 counts in 3 samples. DEseq2 normalized count was used to input of GSEA. MSigDB Hallmark gene set, KRAS-ERK signature^15^, and PDAC subtype signatures were used^2,3,32,62,63^. Single cell RNA-seq derived signatures were only applied to scRNA-seq data. Bulk RNA-seq counts data of pre-post paired samples from WOOM cohort (n=2 for BR, n=5 for non-BR, respectively) or aggregated scRNA-seq count of paired samples (n=2 for BR, n=5 for non-BR, respectively) were extracted and filtered to retain the genes which have more than 10 counts in 30% of samples. DEseq2 analysis was performed with including patient, treatment status, and response status as a covariate in design. The results were ranked by -log10(p-value) *(log2Foldchange) metric and GSEA pre-rank was run. MSigDB Hallmark gene set, KRAS-ERK signature, PDAC subtype signatures were used as well.

### Patienths Single-cell RNA sequencing library construction

Single-cell suspensions were processed according to the 10xGenomics scRNAseq sample preparation protocol (Chromium Next GEM Single Cell 3’ Kit v3.1, 10xGenomics). The entire mixed cell population was further analyzed without sorting or enrichment for specific cell subtypes. Cell suspensions were uploaded into the Chromium controller, capturing GEMs that encapsulated an estimated 5,000-10,000 single cells per channel. Libraries were constructed from the amplified cDNA, and sequencing was performed on the Illumina NovaSeq 6000 platform. All steps were performed according to the manufacturer’s standard protocol.

### Mouse PDAC Single-cell RNA sequencing preparation

Tumors were minced and washed three times with PBS. Small minced pieces were then enzymatically digested for 50 min. with 2.0 mg/ml collagenase A, 1.0 mg/ml Hyaluronidase, and 50 U/ml DNase I in serum-free DMEM at 37 ℃ using continuous mixing conditions. Single-cell suspensions from tumor digestion were prepared by passing tissue through 40-mm nylon strainers. Cells are fixed using Parse Evercode fixation v2 kit and barcoded libraries were made using the Parse Evercode WT v2 kit (Parse Bioscience, Seattle WA) and sequenced by Illumina NovaSeq. Fastq files are made using pipeline provided by Parse Bioscience.

### Human scRNA-seq data processing

The UMI gene count matrix was converted to Seurat Object format using Seurat (v5.1.0). Only cells with greater than 500 unique genes and greater than 750 UMI expressed are retained and integrated by harmony to initial clustering. Doublets were identified and removed within each library using the R package scDblFinder. Computational decontamination was performed by the R package DecontX. Tumor and non-tumor cells containing less than 20% and 10% mitochondrial RNA, respectively, were retained for analysis. UMI counts were log normalized and scaled without centering using Seurat (v5.1.0). The top 2000 variable features were identified using the VST method and harmony integration was performed on the Seurat object. UMAP visualization was performed using the resultant 50 PCA factors. Nearest-neighbor was calculated using the SNN method, unsupervised clustering was performed using the Louvain algorithm, UMAP visualization was performed using harmony reduction. For cancer cell analysis, the public data set was obtained from the human PDAC atlas paper^31^. Only epithelial cell population was used and single nuclear RNA-seq data were removed. Cells with greater than 500 unique genes and greater than 750 UMI expressed were retained and normal epithelial clusters were removed based on UMAP. Then retained data was used as the background dataset and integrated with our scRNA-seq cancer cells.

### Mouse sc-RNA-seq data processing

Filtered UMI gene count matrix was converted to Seurat Object format using Seurat (v5.1.0). Only cells with greater than 200 unique genes, and less than 10% mitochondrial RNA expressed are retained and integrated by harmony to initial clustering. Doublets were identified and removed within each sample using the R package DoubletFinder, respectively. UMI counts were log normalized and scaled without centering using Seurat (v5.1.0). The top 2000 variable features were identified using the VST method and harmony integration was performed on the Seurat object. UMAP visualization was performed using the resultant 50 PCA factors. Nearest-neighbor was calculated using the SNN method, unsupervised clustering was performed using the Louvain algorithm, UMAP visualization was performed using harmony reduction. InferCNV was performed for inferring CNV from scRNA-seq data towards each sample. Subcluster mode with random tree was used and BayesMaxPNormal=0.3 was used for filtering out low probability CNVs.

### scRNA-seq differential expression and gene signature score

Addmodule score function was used for calculating gene signature scores of tumor cell fractions. Since Addmodule score is the relative score in the dataset, we combined PDAC atlas tumor cell dataset with our WOOM tumor cell dataset as a background to assess the relative positioning of cells in the overall PDAC. Sample level mean of the score was compared to find differentially expressed pathways between BR and non-BR.

Cluster based Differential expression analysis was performed using FindMarker function of Seurat and MAST method.

### Ethics for Animal Studies

All animal studies, including the surgical implantation of PDX tissues and KMC /KPC tumor cell lines, were conducted in compliance with IACUC guidelines and in accordance with OHSU protocol No. TR01_IP00001014.

### *In vivo* PDX tumor model

Male NOD.Cg-Prkdcscid Il2rgtm1Wjl/SzJ mice (CAT#5557, The Jackson Laboratory, 10-12 weeks old) were used for transplanting PDX tissues. While under aneshesia, a small incision was made at the shoulder of each mouse, and approximately 5mm tumor chunk was implanted after dipping Matrigel. The incision was closed using surgical clips, and mice were monitored until recovery from anesthesia. Tumor growth was measured by nogis twice weekly, and tumor volume was calculated using the formula: V = (short axis* short axis * long axis)/2. Tumors were allowed to grow until they reached 5mm size of short axis. Once the tumors reached the desired size, they were randomized for experimental use. Mice were taken down when they became moribund.

### *In vivo* orthotopic mouse tumor model

Male B6129SF1/J mice (CAT#101043, The Jackson Laboratory, 10-12 weeks old) were used for transplanting KMC cell lines and female B57BL/6J mice (CAT#664, The Jackson Laboratory, 10-12 weeks old) were used for transplanting KPC cell lines. While under anesthesia, small incision was made at the left side of the abdomen of each mouse, and 5000 cells in 20μl of 50% DMEM/ 50% Matrigel medium were orthotopically implanted into pancreatic tail. The incision was closed using sutures and surgical clips, and mice were monitored until recovery from anesthesia. Tumor growth was monitored twice weekly using ultrasound (Vevo2000, Fuji Film), and tumor volume was calculated using the formula: V = (axial area*supero-inferior length + sag area*right-left length)/3. Tumors were allowed to grow until they reached an average volume of approximately 100 mm³. Once the tumors reached the desired volume, they were randomized for experimental use. Mice were taken down 10 days after treatment for scRNA-seq or when they became moribund.

### DepMap analysis

RNA-seq data, KRAS mutation status, CNV information and drug sensitivity for Cobimetinib, MRTX1133, Gemcitabine, Fuluolouracil, Oxaliplatin, Paclitaxel, Dosetaxel, Irinotecan, Leucovolin, and Erlotinib were obtained from DepMap PRISM_Repurposing_Public_24Q2. Expression data was obtained from Batch corrected Expression Public 24Q4 and GSVA was performed to obtain BR score in KRAS mutated cell lines or *KRAS^G12D^* mutated cell lines with response data for any drugs described above. For CNV analysis, Copy Number Public 24Q4 (Log2 transformed) data was obtained and relative CN was calculated from following formula. (Log2 transformed value) = Log2((relative CN)/2 +1). Then, relative CN>2.5 was called as copy number gain. Drug sensitivity data was obtained from PRISM Repurposing Public 24Q2 and published paper^64^.

### *In vitro* drug treatment

Cells were plated in 96-well, black-walled, clear-bottom plates (Corning, 3914) for cell viability assay (2000 cells of 301J and PDCLs, and 500 cells of 303J). The day after plating, serial dilutions of MRTX1133 (four-fold) and OMO-103 (two-fold) were added to evaluate a concentration-dependent effect on cell viability. Seventy-two hours later, Prestoblue-HS (A13261, Thermo Fisher) reagent was added, and fluorescence was measured after 30 min incubation by Cytation 5 (Bio Tek). GR values were calculated and used for synergy assessment by SynergyFinder. ZIP method was used.

### Statistics

No randomization was performed in our study as it is single arm phase 1 trial. Blinding was not used in any aspect of our study except during histological data appraisal by pathologists, who were blinded to study cohorts. A log-rank test was used to compare K–M survival and recurrence curves as indicated in figures. CPH modeling was used to estimate HRs for survival and recurrence with associated P values. For all survival analysis, only patients alive 30 days or more after surgery were included to avoid analyzing death related to surgical complications. Two-tailed t-tests were used when comparing two conditions and analysis of variance (ANOVA) was used when comparing more than two conditions within a dataset. For non-Gaussian data we used Wilcoxon test. Pearson correlation coefficients were generated for Gaussian data. Two-sided Fisher’s exact tests were used for categorical comparisons. FDR multiple comparisons correction was applied using the Benjamini–Hochberg method.

## Acknowledgemnts

We thank to all the patients enrolled this WOO trial. We thank to Katelyn T. Byrne (OHSU) for sharing 6499c3 and 6499c4 KPC cell lines. This study was supported by the National Cancer Institute (NCI) of the National Institutes of Health under awards; R01 CA186241(to R.C.S.), R01 CA287672 (to J.R.B.), R01 CA212600 (to J.R.B.), R21 CA263996 (to R.C.S. and J.R.B), U01 CA224012 (to R.C.S., L.M.C., and J.R.B), U01 CA294548 (to R.C.S., L.M.C., and J.R.B), U01 CA278923 (to R.C.S.), U01 CA 281902 (to G.B.M), P30 CA069533 (to L.M.C),. This study was also supported by DoD PA210068 (R.C.S., P.J.W, and L.M.C.), DoD Breast Cancer Research Program (L.M.C.), the Susan G Komen Foundation (SAC200100, SAC240100), National Foundation for Cancer Research (L.M.C.), the Knight Cancer Institute (L.M.C.), Hildegard Lamfrom Endowed Chair in Basic Research (L.M.C.), and from the Brenden-Colson Center for Pancreatic Care (R.C.S, G.B.M, L.M.C., J.R.B.)

## Declaration of interests

L.S. is founder, shareholder and employee of Peptomyc SL. L.M.C. has received reagent support from Cell Signaling Technologies, Plexxikon, Inc., and Syndax Pharmaceuticals, Inc.; is on advisory board for Carisma Therapeutics, Inc., CytomX Therapeutics, Inc., Kineta, Inc., Hibercell, Inc., Cell Signaling Technologies, Inc., Alkermes, Inc., NextCure, Guardian Bio, Dispatch Biotherapeutics, AstraZeneca Partner of Choice Network (OHSU Site Leader), Genenta Sciences, Pio Therapeutics Pty Ltd., and Lustgarten Foundation for Pancreatic Cancer Research Therapeutics Working Group, Inc. J.R.B. declares SAB for Perthera (Precision Medicine company), PanCAN (pancreatic cancer advocacy group); co-owner of Faster Better Media (electrophoresis company); and is a chief editor for Taylor & Francis. C.D.L. has received research support from AstraZeneca, Servier, and Genentech/Roche. G.B.M. is on the SAB or consultant: Amphista, Astex, Astrazeneca, Bluedot, Ellipses pharmaceuticals, Immunomet, Leapfrog Bio, Bruker/Nanostring, Neophore, Nerviano, Nuvectis, Pangea, PDX Pharmaceuticals, Qureator, Rybodyne, Signalchem Lifesciences, Tarveda, Turbine, Zentalis Pharmaceuticals, has stock/options/financial: Bluedot, Catena Pharmaceuticals, Immunomet, Nuvectis, Rybodyne, Signalchem Lifesciences, Tarveda, Turbine, Licensed technology: HRD assay to myriad genetics, DSP patents with Nanostring and Sponsored research: Astrazeneca, Zentalis, and Nanostring. R.C.S. has received research funding support from Cardiff Oncology and AstraZeneca. She was a consultant for Rappta Therapeutics. She is currently a consultant for Revolution Medicine, and she is on the scientific advisory board for PanCAN, MOHCCN and PanCuRx Canada. She is an External advisor for Cedars Sinai and on the executive board for PRECEDE.

---

**Figure S1.**
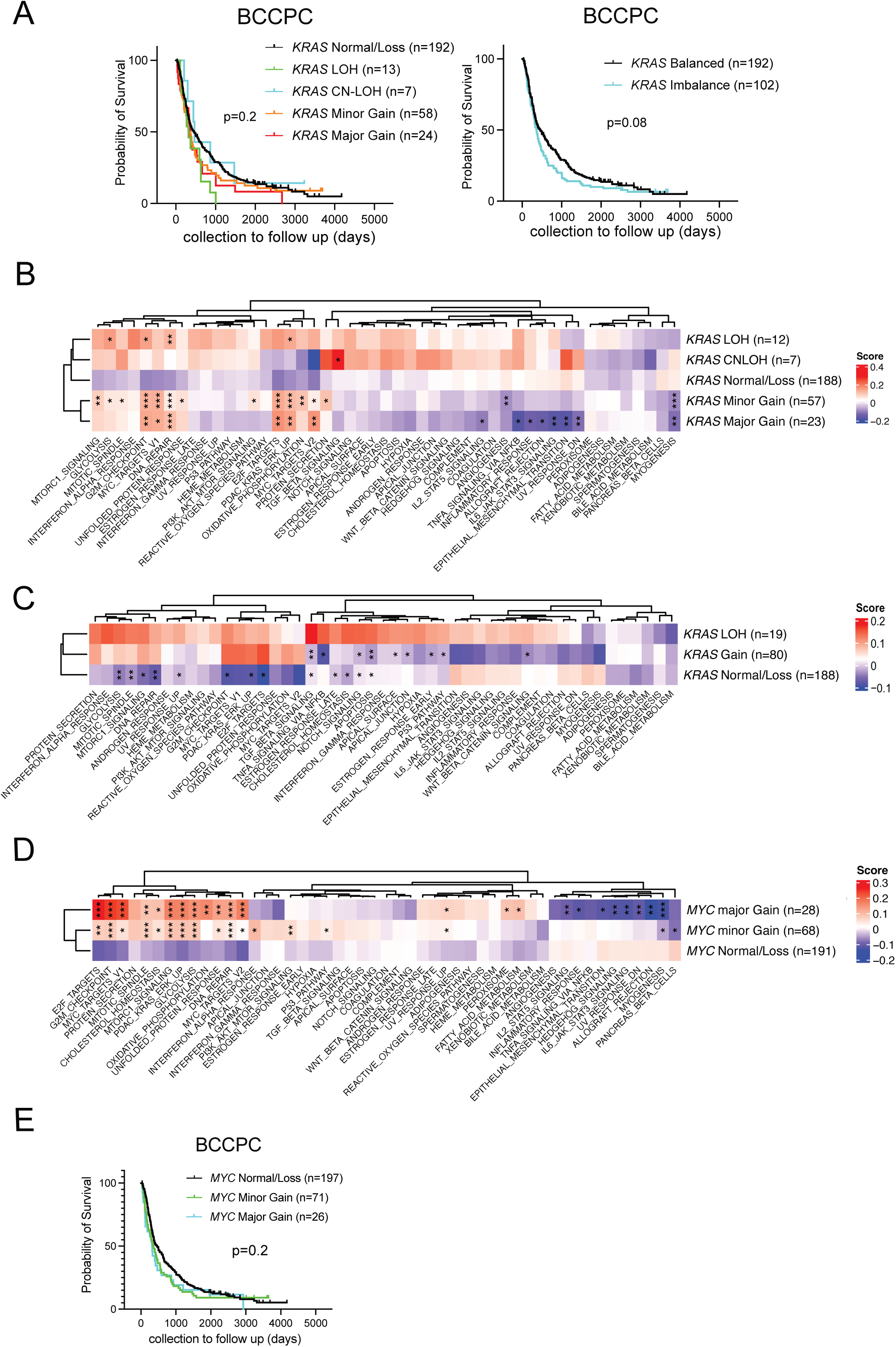
(**A**) Kaplan-Meyer (KM) survival plots of Brenden-Colson Center for Pancreatic Care (BCCPC) cohort (n=294, patients with CNV data, the patients died within 30 days after collection were excluded) stratified by *KRAS* copy number variance (CNV). (Left): *KRAS* copy number was divided into 5 groups. Normal/Loss: CN-neutral and CN=1 without mutation. LOH (loss of heterozygosity): 1 copy loss in *KRAS* mutant patients. CN-LOH (copy neutral LOH): Copy number neutral with both alleles mutant. Suggesting doubling after LOH. Minor Gain: copy number=3. Major Gain: copy number≥4. (Right): *KRAS* copy number status were combined into 2 states. Balanced: Normal/Loss. Imbalanced: LOH, CNLOH, minor Gain and major Gain. Log-rank test. (**B,C**) Heatmaps showing the mean GSVA scores of MSigDB Hallmark and PDAC-KRAS/ERK signature across *KRAS* CNV categories. (n=287, patients with RNA-seq data and CNV data) (**B)** All five groups were compared. (**C)** Gains and LOHs were combined and compared. BCCPC dataset. Dunnet’s test. (Ref: *KRAS* Normal/Loss(**B**) and *KRAS* LOH(**C**)). *:p<0.05, **: p<0.01, ***: p<0.001 (**D**) Heatmap showing mean GSVA score of MSigDB Hallmark and PDAC-KRAS/ERK signature across *MYC* CNV categories. BCCPC cohort (n=287, patients with RNA-seq data and CNV data). Minor Gain: *MYC* CN=3, Major Gain: *MYC* CN>=4. Dunnet test. Reference: *MYC* Normal/Loss. (**E**) KM survival plot of BCCPC cohort (n=294,patients with CNV data, the patients died within 30 days after collection was excluded). Log-rank test.

**Figure S2.**
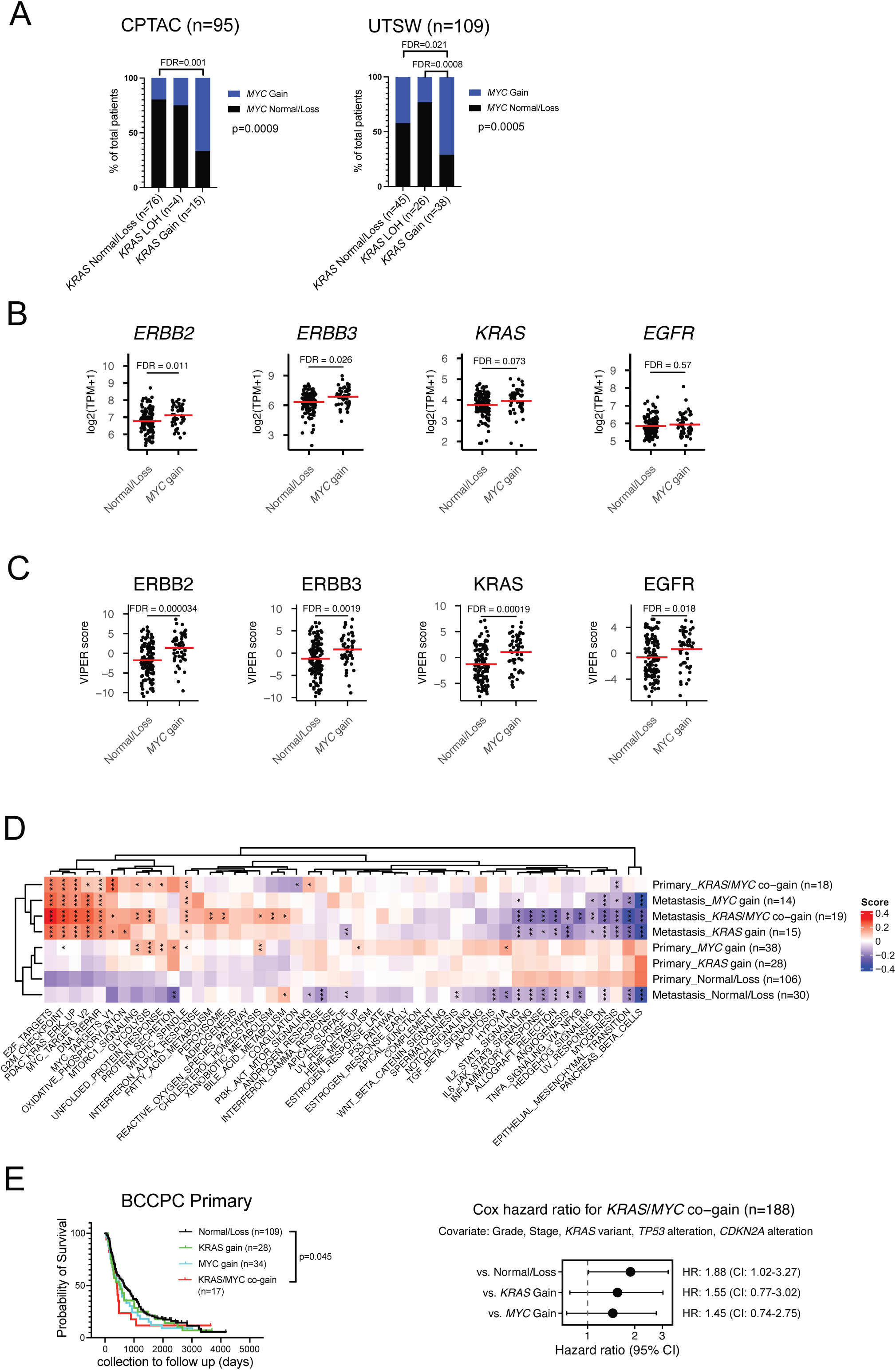
(**A**) Relationship between *KRAS* CNV and *MYC* CNV in CPTAC (left; n=95 with CNV) and UTSW (right; n=109 with CNV) cohorts. p value describes whole group comparison by Fisher’s exact test. Pairwise comparisons were performed using the Wilcoxon test, with FDR correction applied across all pairwise tests. (**B**) Gene expressions of KRAS upstream ERBB signalings (*EGFR*, *ERBB2*, and *ERBB3*) and *KRAS,* compared between *MYC* Gain (n=52) and Normal/Loss (n=136). BCCPC cohort (n=188; *KRAS* Normal/Loss patients only). Wilcoxon test. FDR corrected with all genes. (**C**) Predicted protein activity of RAS upstream ERRB signalings (*EGFR*, *ERBB2*, and *ERBB3*) and KRAS by VIPER in primary PDAC between *MYC* Gain (n=52) and Normal/Loss (n=136). (n=188; *KRAS* Normal/Loss patients only). VIPER scores were derived fromour previous publication (Link JM, *et al.* Nat Cancer. 2025). Wilcoxon test. FDR corrected with all genes. (**D**) Heatmap of mean GSVA scores of MSigDB Hallmark and PDAC-KRAS/ERK signature across *KRAS* and *MYC* copy number status in primary and metastatic PDAC, respectively, in BCCPC cohort (n=268, with CNV and RNA-seq data, *KRAS* LOH was excluded.). (Ref: Primary_Normal/Loss). One-way ANOVA, Dunnet test. *: p<0.05, **: p<0.01, ***: p<0.001 (**E**) KM survival plot and forest plot regarding *KRAS* and *MYC* gain status in primary PDAC of BCCPC cohort (n=188; with CNV data, *KRAS* LOH and patients died within 30 days of enrollment were excluded.). Normal/Loss/Gain only. Cox Hazard Multivariate analysis. Adjusted by the same factors used in whole dataset (Fig. 1F) plus lymphovascular invasion (significant in Univariable analysis). P value is from Cox Hazard Multivariate analysis.

**Figure S3.**
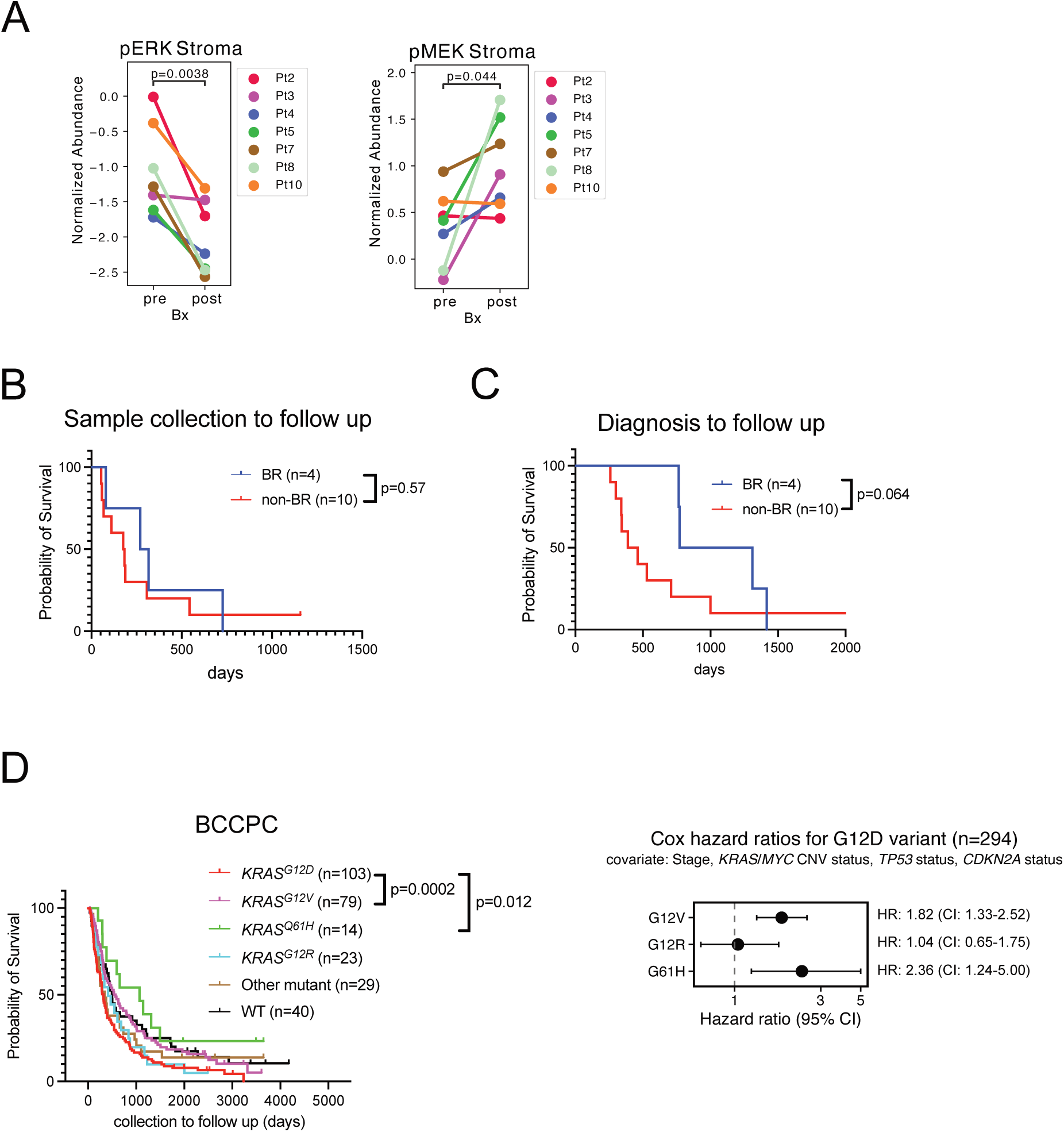
(**A**) Digital Spatial Profiling (DSP). Comparison of normalized protein abundance of tumor adjacent normal area between pre- and post-treatment. Phospho-ERK (Left; n=7) and phospho-MEK (Right; n=7). (**B**) KM survival plot for sample collection to follow up between BR (n=4) and non-BR (n=10). Log rank test. (**C**) KM survival plot for diagnosis to follow up between BR (n=4) and non-BR (n=10). Wilcoxon test. (**D**) KM survival plot and forest plot regarding *KRAS* mutant variants in BCCPC dataset (n=294; with RNA-seq data and CNV dta). Cox Hazard Multivariate analysis adjusted by tumor stage, *KRAS*/*MYC* CNV status, and mutation status of *TP53* and *CDKN2A*. P value in KM plot is from Cox Hazard Multivariate analysis.

**Figure S4.**
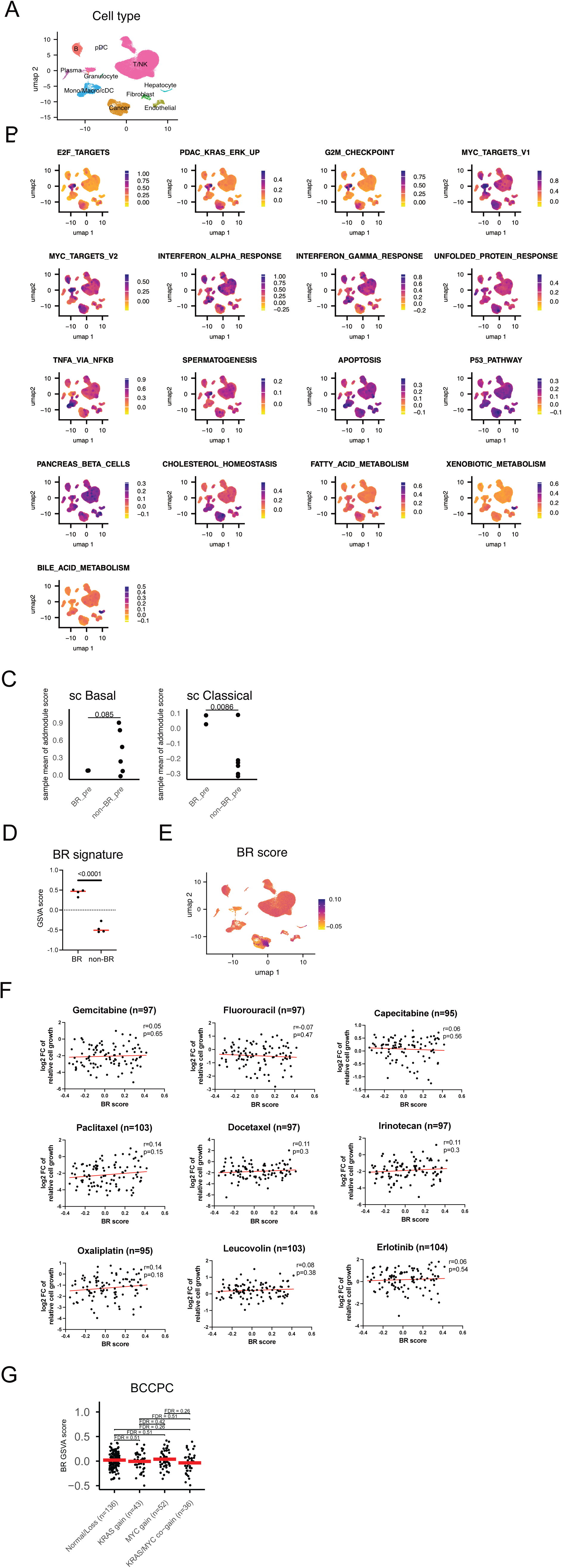
(**A**) UMAP of WOOM single cell RNA-seq (n=7 patients) colored by cell types. (**B**) The scores of differentially expressed pathways between BR-pre and nonBR-pre bulk RNA-seq (Showing in Fig. 3A) were mapped onto UMAP of WOOM single cell RNA-seq. AddmoduleScore. (**C**) Graphs showing subtype differences between BR (n=2) and nonBR (n=5) samples in tumor cells of scRNA-seq. Sample level mean values of AddmoduleScore. (**D**) BR GSVA score in BR-pre (n=4) and nonBR-pre (n=4) patients. RNA-seq. (**E**) Feature Plot of BR score. scRNA-seq. AddmoduleScore**. F**: Correlation of BR GSVA score and *in vitro* response to the PDAC standard of chemotherapy drugs on DepMap. Pearson. **G**: Correlation between BR GSVA score and *KRAS*/*MYC* gain status. BCCPC dataset(n=267; with CNV and RNA-seq, KRAS LOH was excluded).

**Figure S5.**
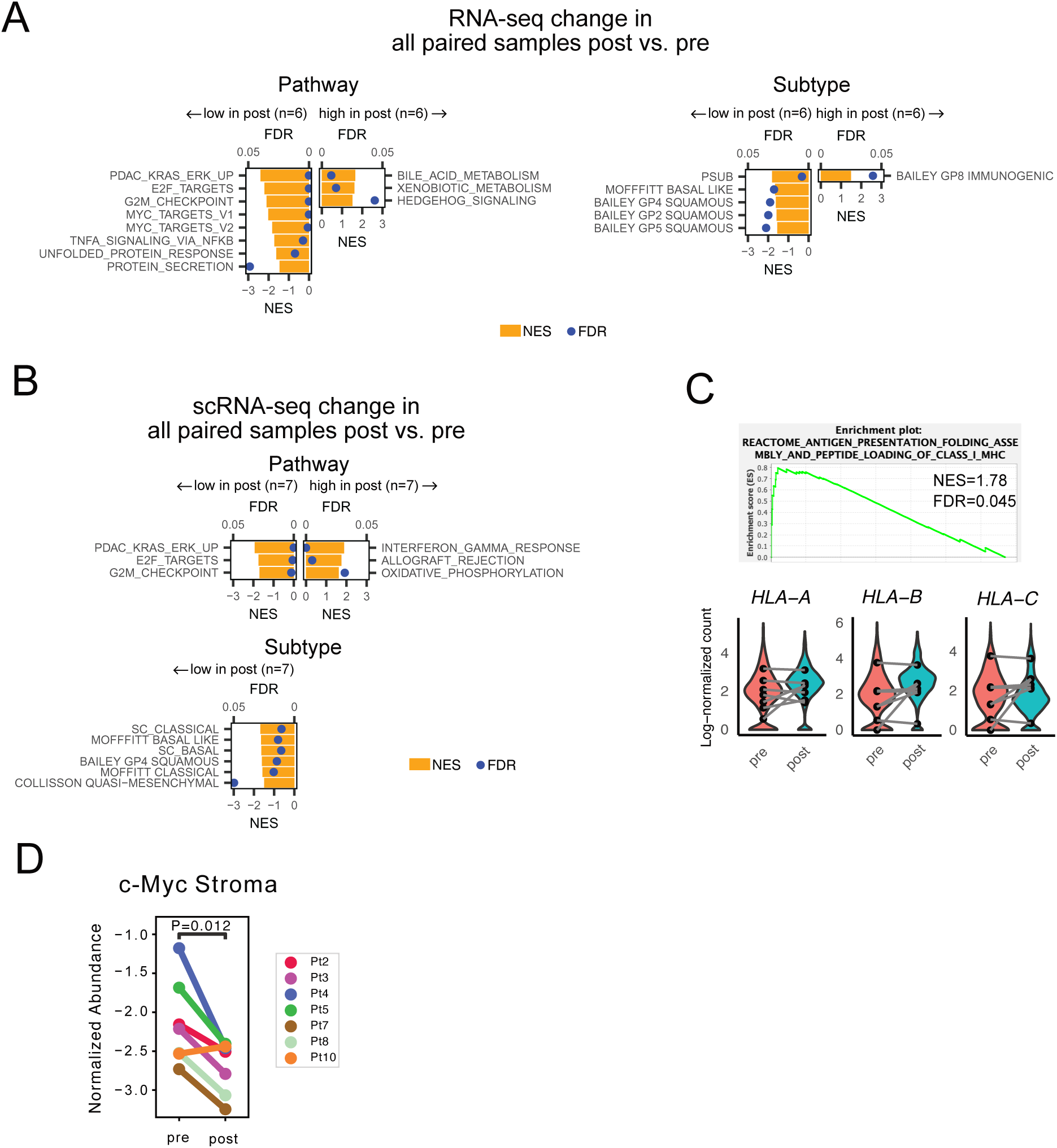
(**A**) Bulk RNA-seq. GSEA prerank of DEGs between post- and pre-treatment in paired patients(n=6). Ranked metric: log10(p-value)*Log2FoldChange. DESeq2. (**B**) Pseudobulk scRNA-seq. GSEA prerank of DEGs(DEG) between post- and pre-treatment in paired patients(n=7) Ranked metric: log10(p-value)*Log2FoldChange. DESeq2. (**C**) (Top): Pseudobulk scRNA-seq. GSEA prerank of MSigDB C2 CP Reactome showing activation of the “Antigen Presentation: Folding, assembly and peptide loading of class I MHC” pathway in post-treatment tumor. Ranked metric: log10(p-value)*Log2FoldChange. DESeq2. (n=7) (Bottom): Violin plot showing HLA class I gene texpression level between pre and post samples (n=7). Violin plots represent the cell-level distribution and dots show the sample-level mean scores. (**D**) DSP. Comparison of normalized protein abundance of tumor adjacent normal area between pre- and post-treatment samples for c-MYC (n=7).

**Figure S6.**
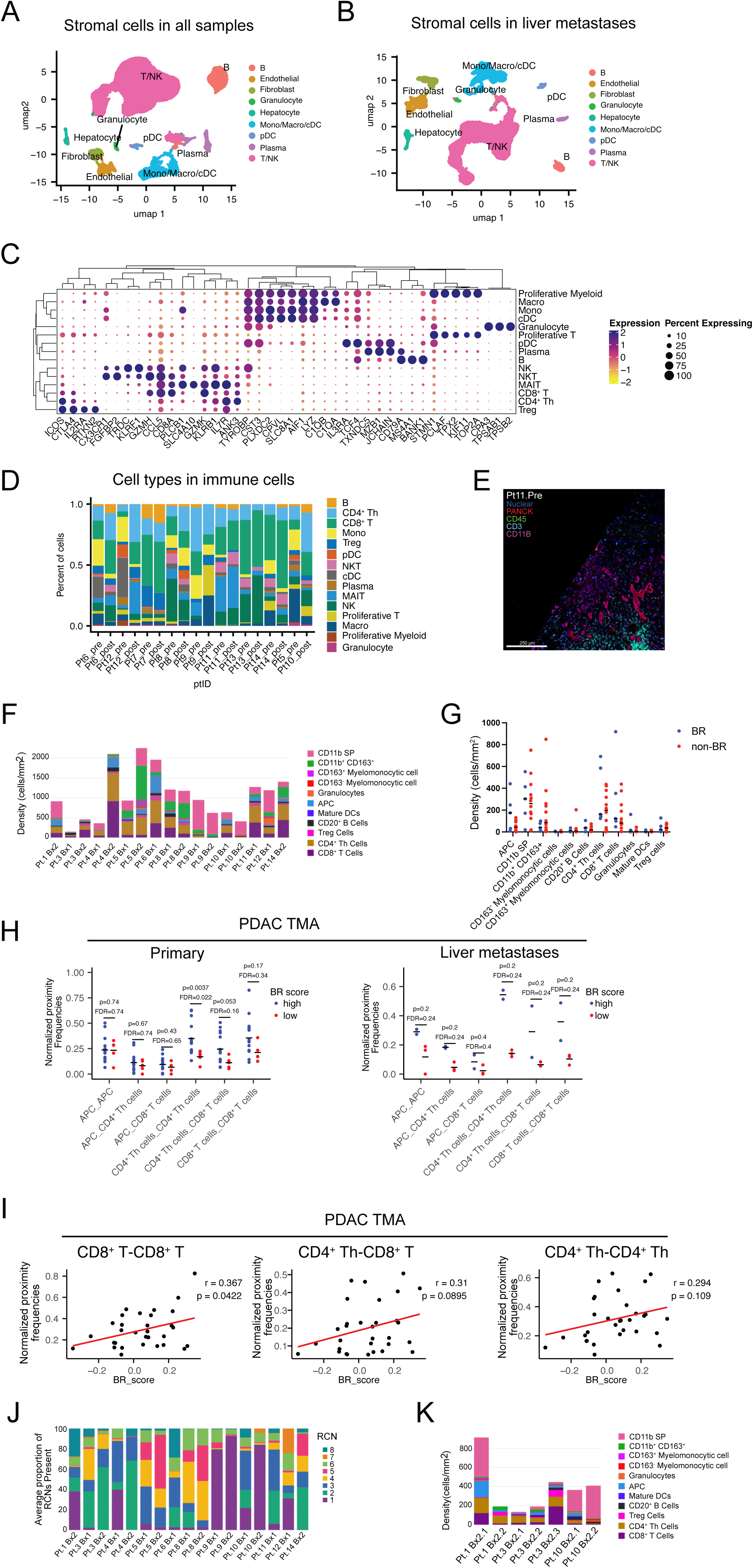
(**A**) UMAP of stromal cell populations from WOOM scRNA-seq (n=7; 6 liver metastases and 1 lymph node metastasis). (**B**) UMAP of stromal cell populations of from WOOM scRNA-seq restricted to liver metastases samples (n=6). (**C**) Enriched marker genes for detailed immune cell types in WOOM scRNA-seq. Top 3 marker genes per cell type were selected using FindAllMarker function of Seurat. (**D**) Fraction of detailed immune cell types towards individual samples in WOOM scRNA-seq. Pt.6 and Pt.12 represent BR tumors. (**E**) Representative pseudo-color images of major cell lineages in WOOM mIHC. (**F**) Density of each Immune cell type per sample quantified by WOOM mIHC. Pt.1,4,6,11 represent BR tumors. (**G**) Dot plot comparing immune cell densities in WOOM mIHC between BR (n=5) and non-BR (n=12). Each dot represent sample-level mean of ROIs. (**H**) Immune cell proximity of PDAC TMA mIHC between BR high and low. (Left)Primary (n=20) and (Right) liver metastases (n=5). Proximity scores were normalized by cell density. (**I**) Scatter Plots of BR_score versus T cell-T cell proximity scores in PDAC TMA mIHC (n=31). (Left)CD8^+^-CD8^+^ T, (Middle)CD4^+^ Th-CD8^+^ T, (Right)CD4^+^ Th-CD4^+^ Th T. (**J**) Percentage of each neighborhood of each sample in WOOM mIHC. Pt.1,4,6,11 are BR tumors. Cellular neiborhood in 60μm radius. (**K**) Density of each Immune cell type in technical duplicate samples measured by WOOM mIHC.

**Figure S7.**
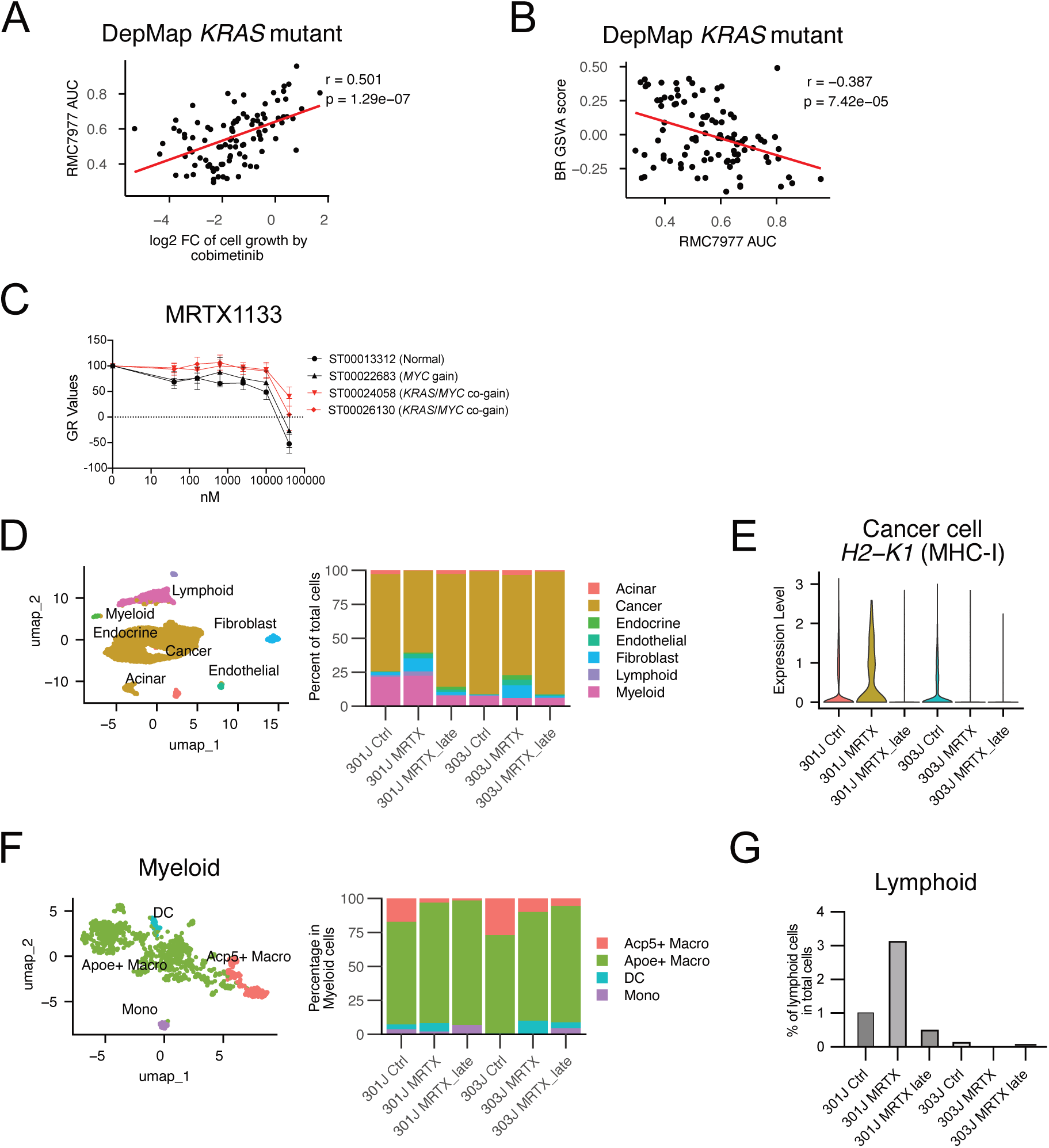
(**A**) Correlation between cobimetinib response and RMC7977 response (area under the curve: AUC) in *KRAS* mutated cell lines (n=99). DepMap. (**B**) Correlation between BR signature and RMC7977 AUC in *KRAS* mutated cell lines (n=138). DepMap. (**C**) Dose response curve for MRTX1133 treated *KRAS^G12D^* mutated low passage patient derived cell lines. *KRAS* and *MYC* CNV status of parental PDAC is shown next to the each cell line name. (**D**) UMAP and proportions of each cell types from scRNA-seq of orthotopic tumors of KMC301J and KMC303J. (**E**) Expression level of *H2-K1* gene in cancer cells. scRNA-seq. (**F**) UMAP and proportions of each detailed myeloid cell type from scRNA-seq of orthotopic tumors of KMC301J and KMC303J. (**G**) Percentage of lymphoid cells among total cells across treatment conditions. scRNA-seq.

